# Effect of differential protection regimes on the diversity and composition of woody plants in the Western Ghats

**DOI:** 10.1101/2024.09.27.615373

**Authors:** Bhushan K. Shigwan, Aboli Kulkarni, Vijayan Smirthy, Navendu V. Page, Rohan Shetti, Mandar N. Datar

## Abstract

Human interference in forests is inevitable, and despite significant conservation efforts, many forest areas continue to suffer from anthropogenic pressures. The forests of the Northern Western Ghats (NWG) exhibit varying degrees of protection, including private, community, and legal frameworks. However, the tree diversity within these protection regimes remains underexplored. This study aims to assess tree diversity, composition, and structure across four protection regimes using a transect-cum-quadrat method, with four quadrats (20×20 m) along a single transect line. Data collected included species richness, individual counts, girth at breast height (GBH), basal area, and a combined disturbance index (CDI). Approximately 50% commonality was observed among sites across the four protection regimes. Protected Areas (PA) and Reserved Forests (RF) exhibited higher tree densities compared to Sacred Groves (SG) and Private Forests (PV). Notably, Sacred Groves, despite experiencing high disturbance levels, displayed a similar tree variety to PAs and RFs. While species composition across the four regimes was comparable, structural elements such as tree density and basal area varied significantly. Sacred Groves were predominantly characterized by older trees, whereas RFs and PAs were primarily populated by younger trees. These findings underscore the critical need for targeted conservation strategies that address the unique challenges faced by each protection regime. Enhanced conservation planning is essential to mitigate the impacts of disturbances, such as climate change and land use changes, which threaten the biodiversity of these forests. The study highlights the importance of preserving Sacred Groves and emphasizes the role of community involvement in conservation efforts to safeguard endemic species and maintain ecological balance in the NWG.

## 1.1 Introduction

The formal protected area network which includes Reserve Forest, Wildlife Sanctuaries and National Parks, aim to limit anthropogenic impact on ecosystems, safeguard biodiversity, reduce deforestation, and mitigate climate change globally by preserving critical habitats and ecosystem services (Chape et al., 2005; Janishevski et al., 2015). In addition to these legally protected areas, various conservation initiatives such as traditionally conserved forests (sacred groves) and community-conserved forests (Nautiyal & Kaechele, 2007; Berkes, 2009) play a significant role in global conservation management. Similarly, privately owned forests also represent one of the conservation categories where human interference may be completely or partially restricted. These traditional to modern conservation efforts have been individually assessed for their effectiveness by evaluating the biodiversity they support (Chape et al., 2005; Gardner et al., 2007; Shahabuddin & Rao, 2010).

Building on the understanding of various conservation strategies, Shahabuddin and Rao (2010) compared the floristic and faunal components of community-conserved areas with those of formally protected areas, revealing that community-conserved areas were more effective in conservation than strictly protected areas.”

India possesses a profound history of forest conservation, characterized by a strong integration of cultural values, governmental responsibilities, and a longstanding tradition of nature worship (Chaturvedi, 1992; Chandran, 1997; Malhotra et al., 2001; Roy & Fleischman, 2022). Human intervention in the forest is unavoidable as large proportion of population residing in and around the forests is dependent on forests for their livelihood. Despite various conservation efforts mentioned above, human intervention affects the forest over the period and alters forests’ species composition and biodiversity. Several studies have been conducted to check the effectiveness of conservation and influence of anthropogenic pressure in terms of biodiversity richness, compositional changes, biomass and carbon content variation, forest cover depletion, socio-economical influences on state-protected areas, reserve forest, community-protected areas and non-protected or private forests from various region of India (Khan et al., 1987; Bhagwat et al., 2005a,b; Nautiyal & Kaechele, 2007; Ambinakudige & Sathish, 2009; Page et al., 2010; Bdoor, 2016; Ghosh-Harihar et al., 2019; Pradhan et al., 2017, 2019; Rath et al., 2020).

The Western Ghats of India, recognized as a biodiversity hotspot, exemplifies the intricate relationship between high levels of endemism and the alarming loss of primary forest cover, which has been exacerbated by anthropogenic pressures over time (Chandran, 1997). Various forest protection regimes can be observed in the region based on the function and purpose of forests. It has a fairly high density of sacred groves, legally protected areas like reserve forests and protected areas, and private forests which are conserved for multiple purposes like coffee plantation or silviculture and livelihood (Bhagwat et al., 2005a,b; Ambinakudige and Sathish, 2009; Page et al., 2010). Researchers have studied different regimes to analyse forest composition, diversity and distribution and the influencing aspects such as abiotic, biotic and anthropogenic activity and socio-economical value individually. A few studies from the central Western Ghats have attempted to understand the forests-human interaction from sacred groves, state reserve forests and coffee plantations and their impact on biological components (Bhagwat et al., 2005a,b; Ambinakudige and Sathish, 2009; Page et al., 2010).

The NWG differ from the Central and Southern Western Ghats in terms of climate, geology and geography and disturbances the region faces (Kulkarni et al., 2023). The forests in this area are extensively fragmented and subjected to intense human intervention. While the floristic components of NWG have been extensively documented, there has been comparatively limited research focused on the ecological dynamics of its forest ecosystems (Shigwan et al., 2024a). Several protected areas (Kanade et al., 2008, Joglekar et al., 2015), reserve forests (Tadwalkar et al., 2020) and sacred groves (Kulkarni et al., 2018) have been assessed for tree species composition and diversity and the effect of disturbance on vegetation. However, most ecological studies have been conducted at the local level or as isolated investigations, and there is a notable lack of comprehensive research that compares tree diversity, community structure, and composition across the four protection regimes within the NWG as a cohesive unit. A recent study from the NWG reported the combined effect of water stress and chronic disturbance on vegetation from different regimes with various degrees of disturbances (Biswas et al., 2024). Though they have sampled different protection regimes but could not cover the entire NWG. Moreover, this work aims to understand interactive effect of climate and historic disturbance on community structure and composition and their comparison across these regimes was secondary. Tamhane et al. (2024) conducted a study on dense forests in the crest of the NWG, regardless of protection regimes, and identified elevation and humidity as the most influential factors for vegetation within this short latitudinal range. As such, these regimes must be investigated with a primary focus considering the degree of disturbance separately (Shigwan et al. 2024b).

Therefore, the present work attempts to compare the tree vegetation from four major protection regimes considering them as a proxy of disturbance based on their degree of protection. This study covers diversity, composition and structure spread across four protection regimes: Protected areas (PA), reserve forests (RF), sacred groves (SG) and private forests (PV). This work aims to find out the role of protection regimes in shaping the tree vegetation in the fragmented forests of the NWG.

## 1.2 Methods

### 1.2.1 Sampling methods and Data collection

The vegetation sampling was conducted using the transect cum quadrat method based on the earlier studies from the Central Western Ghats (Chandran & Mesta, 2001; Chandran et al., 2010) which are recognized to be suitable for highly fragmented regions. On a 140m transect line, four quadrats of 20×20m were laid alternately with a 20m interval between each quadrat. Individuals with girth at breast height (GBH) of ≥30 cm were recorded from each quadrat along with their taxonomic identity (Mesta & Hegde, 2018). A total of 40 transects (10 in each protection regime) each having four quadrats were laid in the forest fragments across the selected four protection regimes (Figure 1). A total of 40 selected transects were laid in various fragmented forests of various protection regimes such as Protected Areas, Reserve Forests, Sacred Groves and Private forests (Figure 1). Specifically, ten sites from each protection regime were allocated at various elevations among forty sites. These were spread across NWG elevational (26–1411 m a.s.l.) and latitudinal gradients (15.81°–20.00°N). The sampling was carried out from 2017 to 2021, between the pre-monsoon and post-monsoon periods (December to May). This period was selected for various reasons, including accessibility issues due to heavy monsoon rains, data collection challenges, plant phenological conditions, and restrictions in PA and RF for wildlife conservation during the breeding season. Forest fragments were selected based on the disturbance criteria, accessibility for sampling and the availability of regimes

**Figure 1.**
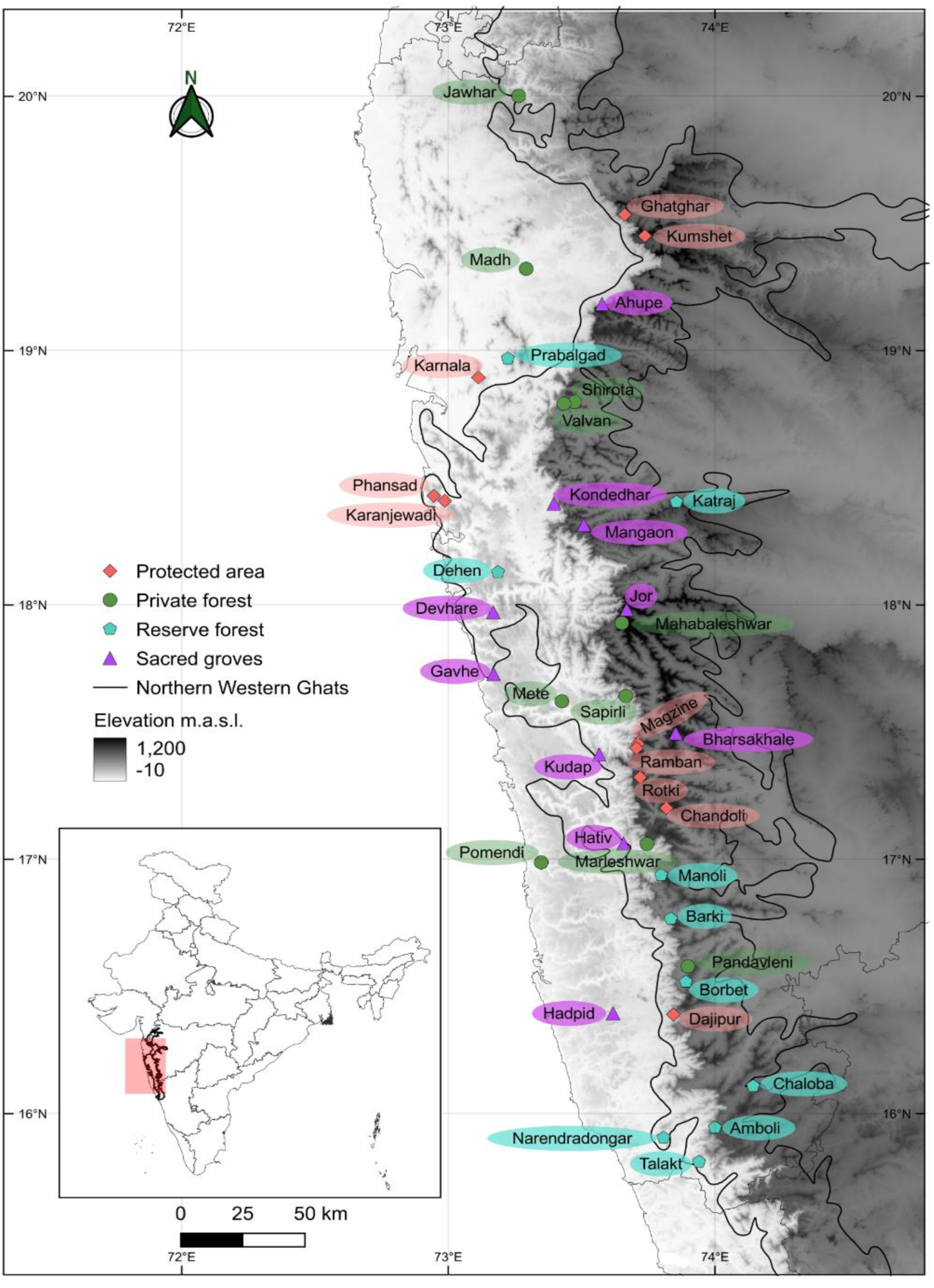
Map depicting transect sites across protection regimes in the NWG

### 1.2.2 Species identification

Species were identified using local flora, field guides (Pascal & Ramesh, 1987; Lakshminarasimhan, 1996; Singh & Kartikeyan, 2000; Singh et al., 2001; Yadav & Sardesai, 2002; Ingalhalikar, 2022), and by comparing them with the specimens from the Agharkar Herbarium of Maharashtra Association (AHMA).

Along with these, elevation and geographical coordinates of each location were also recorded. The disturbances were recorded for eight parameters viz cutting, lopping, construction, fire, grazing, internal pathways, other human activity (e.g. village festivals and rituals, recreational and tourism), and distance from the road from each transect site across protection regimes. An assessment of the disturbance was made by assigning a disturbance score ranging from 0 to 5 for each parameter, based on the observations made during sampling. A score of 0 indicates no evidence of disturbance, while a score of 5 indicates the highest level of disturbance. The total score for each site was then converted into a combined disturbance index (CDI) following Kanade et al., (2008). Additionally, the number of endemic, evergreen and deciduous species, and mode of seed dispersals (Autochory, Anemochory and Zoochory) using secondary literature were also recorded for each protection regime.

### 1.2.3 Data analysis

All the transect data was arranged according to the protection regimes and comparative analysis was carried out.

To understand the distribution of species abundance across the protection regimes, rank abundance curve analysis was carried out. The abundance log_10_ across the rank was plotted using *vegan* and *BiodiversityR* packages in R software (Version 4.2.3, R Core Team, 2023). The goodness of fit was also tested for the models (Geometric, Log series, Broken stick and Log normal) with *chi^2^* and its probability values were obtained using PAST software (Version 4.03). If *the p-value* is greater than 0.05, then the model can be considered to be fit (Magurran, 2004).

The ecological importance of each species was estimated using the Importance Value Index (IVI) (Curtis & McIntosh, 1951; Chandran et al., 2010) across each protection regime with pooled species data. Similarly dominant families were recorded using Family Index Value (FIV) in each regimes following Kanade et al., (2008).

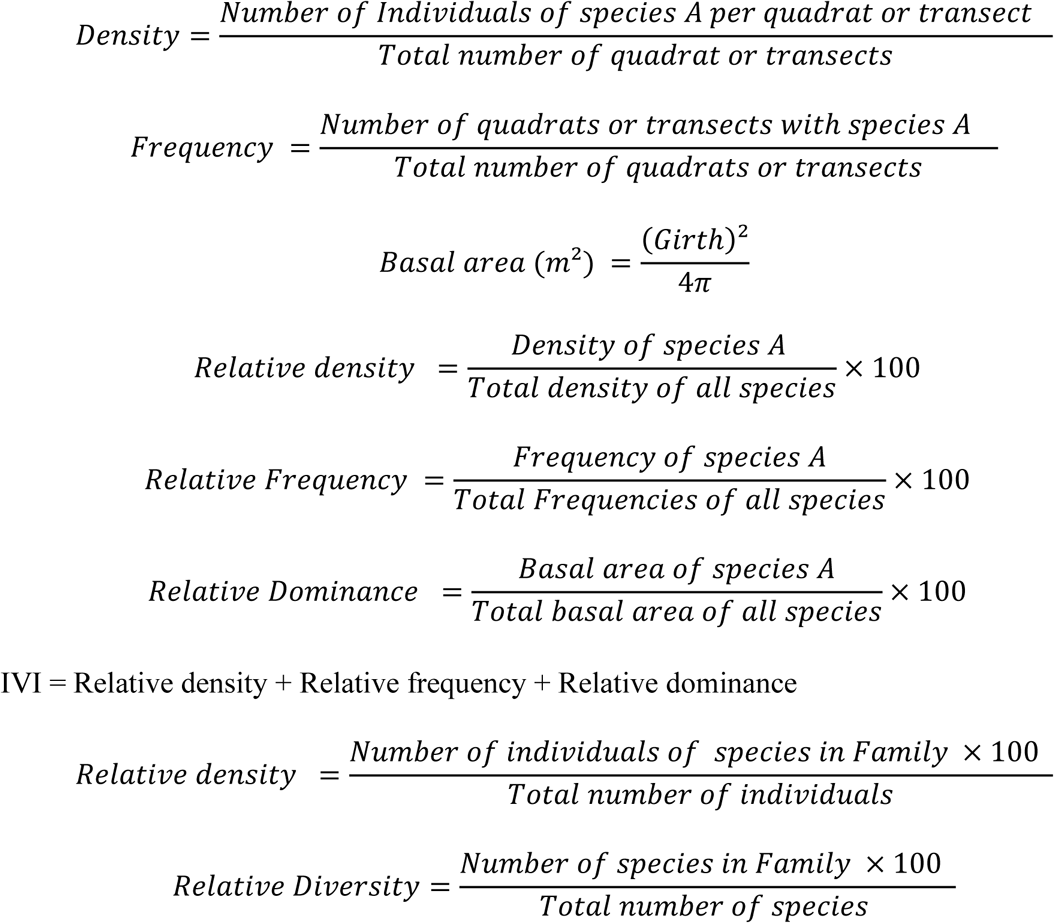

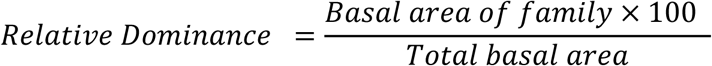

Family Importance Value (FIV) =Relative density + Relative diversity + Relative dominance Richness and diversity: Species richness per transect site (i.e. number of species) was noted. Similarly, Shannon diversity based on abundance per transect site for each protection regime was obtained using PAST software and their means were compared using boxplots.

The percentage of evergreen and deciduous trees along with the mode of seed dispersal was calculated for the transect sites for each protection regime. The structural aspects such as tree density and basal areas were estimated following Chandran et al., (2010) across the regimes. All the vegetation parameters (Richness, Shannon diversity and proportion of Evergreen, Deciduous, Endemics, Autochory, Anemochory and Zoochory) and CDI were compared among all protection regimes using box plots prepared in R software. The comparisons were supported by a non-parametric Kruskal–Wallis test.

To understand the similarity and dissimilarity between the protection regimes, a cluster analysis was performed in the PAST software with pooled data of abundance of each species using the Bray-Curtis method. Further, to visualize and explore the similarities and dissimilarities, NMDS was performed with site-wise abundance data across protection regimes using the Bray-Curtis method in R software (Oksanen, 2022).

The structural aspects (tree density and basal area) per protection regimes were further compared with the box plots and were supported by the Kruskal-Wallis test. Separate investigations were also conducted into the distribution of tree density and basal area by girth classes (30–60, 61–90, 91–120, 121–150, 151–180, and 181–500) by comparing it among four regimes with box plots. The Kruskal-Wallis test was used to determine the statistical significance.

## 1.3 Results

### 1.3.1. Species diversity

The present investigation recorded a total of 3360 individuals under 148 taxa belonging to 43 families and 118 genera. Out of total abundance, only 0.1% of taxa were identified only to the genus level, while rest other to specific level. All of them were included in the analysis. A comparative analysis of various parameters obtained from four protection regimes based on these is provided in Table 1. The overall species richness, genera, number of individuals, evergreen richness, endemics, and endemic individuals recorded from RF were comparatively higher compared to other regimes, according to the species aggregated data from all protection regimes (Table 1). Whereas, PVs were noted with less evergreen (richness and individuals) and high deciduous (richness and its individuals) than other regimes.

**Table 1.**
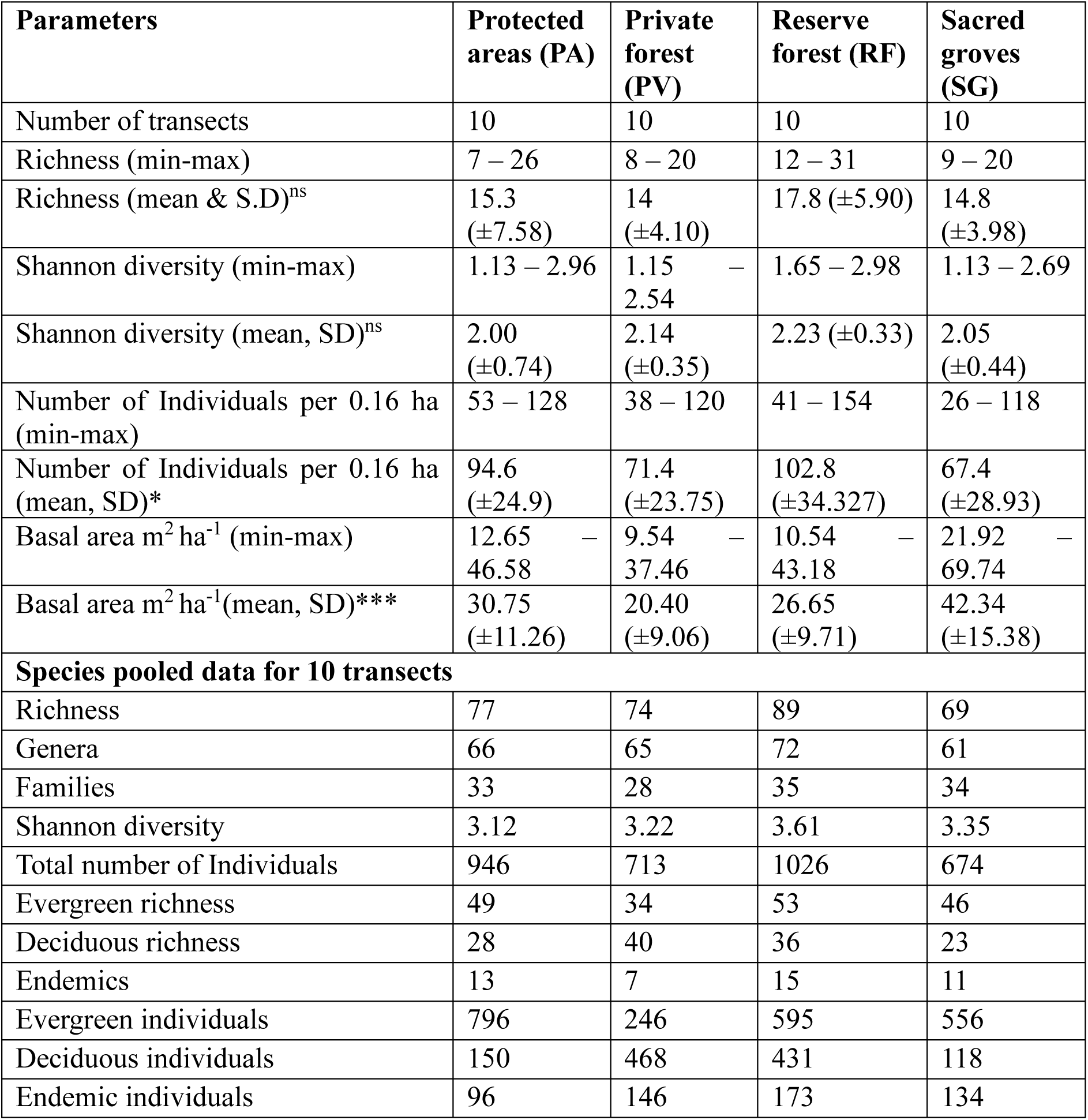
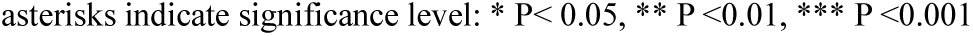
Overview of vegetation parameters across protection regimes obtained from quantitative assessment.

All protection regimes shared only 26 species and showed unique species ranging from 11 to 20 (Figure S1). The family Lauraceae was recorded as species-rich family from PA and RF and second most from SG. Leguminosae was species-rich from PV followed by Rubiaceae and Malvaceae. Whereas SG showed Moraceae as most speciose followed by Lauraceae and Rubiaceae (Table S1).

### 1.3.2 Dominant communities across protection regimes

The analysis of the species abundance in PA resulted in only three species (*Memecylon umbellatum, Olea dioica,* and *Dimocarpus longan*) accounting for around 47% of the total abundance, while the remaining 74 species contributed to the 53% of the abundance. From PV forests, four species (*Terminalia elliptica, T. paniculata, Syzygium cumini* and *M. umbellatum*) contributed 47% of the abundance and 70 species to the remaining 53% of the total abundance. Seven species (*M. umbellatum, T. elliptica, S. cumini, T. paniculata, O. dioica, Atlantia racemosa* and *Xylia xylocarpa*) recorded from RF contributed 47% of abundance and remaining 82 species contributed 53% of total abundance. Similarly, six species (*M. umbellatum, Garcinia talbotii, Ixora brachiata, Mangifera indica, Dimocarpus longan* and *Olea dioica*) contributed 50%, while 63 species contributed to the remaining 50%. *Memecylon umbellatum* was among the abundant species across all types of protection regime.

The rank abundance curves for all four regimes were found to be following the log-normal curve model (Figure S2). This was also supported by the *chi^2^* value and *p* values illustrated in Figure S2 (Table S2).

The ecological importance of each species across the protection regimes was investigated using IVI. Ten species with high IVI per protection regimes are represented in Figure S3. It was observed that *Memecylon umbellatum* (MEMUMB, 52.49), *Olea dioica* (OLEDIO, 35.33) and *Syzygium cumini* (SYZCUM, 14.7) were with the highest IVI from PA. Whereas, SG had highest IVI for *Memecylon umbellatum* (MEMUMB, 32.24), *Mangifera indica* (MANIND, 30.40) and *Syzygium cumini* (SYZCUM, 16). In RF, high IVI was represented by *Ficus nervosa* (FICNER, 26.89), *Syzygium cumini* (SYZCUM, 23.07) and *Memecylon umbellatum* (MEMUMB, 18.86). While *Terminalia elliptica* (TERELL, 37.30), *Terminalia paniculata* (TERPAN, 31.38) and *Syzygium cumini* (SYZCUM, 24.39) were recorded with high IVI from PV (Table S3).

In case of FIV, four different families were found to be dominant in each protection regime (Table S4). The dominant family were recorded such as Melastomataceae (49.87) from PA, Combretaceae (70.05) from PV, Moraceae (38.59) from RF and Anacardiaceae (35.37) from SG.

### 1.3.3 Similarity and dissimilarity across protection regimes

To assess the degree of similarity among the various protection regimes, species abundance data from all ten transect sites within each regime were compiled. The cluster analysis of this data yielded two clusters, SG – PA and PV – RF with a similarity of approximately 38%. While, the cluster SG – PA showed 50% resemblance, PV – RF demonstrated 54% similarity (Figure S4).

The NMDS analysis, which examined site-wise species abundance across protection regimes, indicated that over 25 sites clustered closely together at the center (Figure 2). Only a few sites from the PV category, specifically JAWH, MADH, and SAPI, along with one site from the RF, KTRJ, were not grouped with the others. Additionally, one more site MANO from the RF category was positioned distinctly away from the other sites.

**Figure 2.**
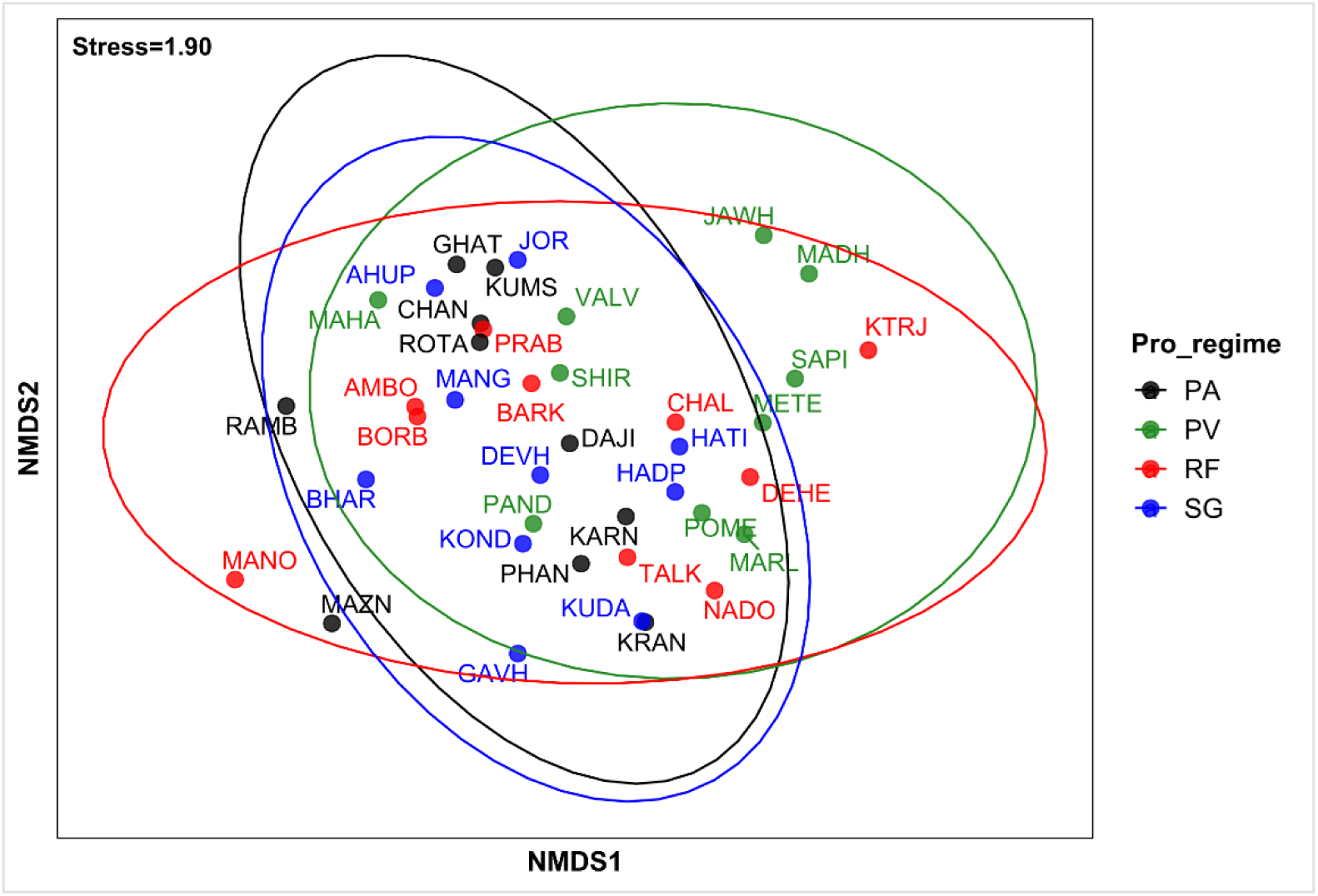
Non-metric Multidimensional Scaling (NMDS) plot comparing species composition across four protection regimes, demonstrating similarities and differences among locations

### 1.3.4 Comparison of disturbance and vegetation parameters

The disturbance factor (CDI) was found to be significantly different across protection regimes. A higher disturbance was recorded from SG and PV than from PA while RF with medium CDI value (Figure 3 A). In the case of species richness and Shannon diversity index (*H*), no significant difference was observed (Figure 3 B-C). Similarly, no significant difference was found in the percentage of evergreen and deciduous species between all four protection regimes (Figure 3 D-E). However, PA and PV sites were found to be significantly different in percentage of evergreen (*p* < 0.05) and deciduous (*p* < 0.05) species. The percentage of species falling under different modes of seed dispersal was also compared between protection regimes (Figure 3 F-H), revealing that the percentage of autochory species was significantly different. Whereas, the percentage of zoochory and anemochory species were not significantly different.

**Figure 3.**
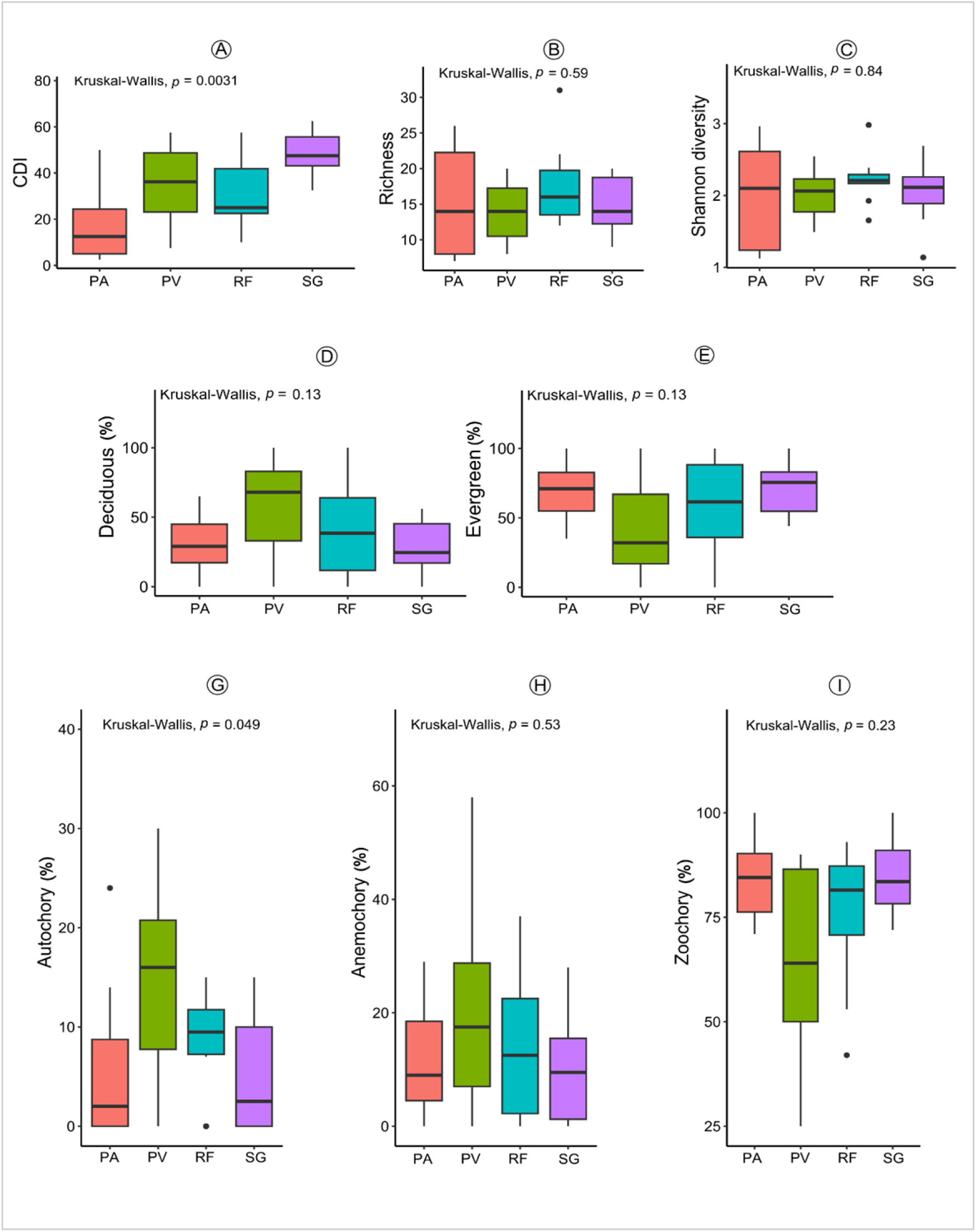
Boxplots comparing disturbance and vegetation parameters among protection regimes. A-Disturbance (CDI); B-E vegetation parameters: B-Richness, C-Shannon diversity index (*H*), D-percentage of Evergreen species, E-percentage of Deciduous species; G-H Seed dispersal modes in percentage: G-Autochory, H-Anemochory, I-Zoochory

### 1.3.5 Community structure across protection regimes

The comparison of structural characteristics of tree communities revealed significant differences in tree density among the various protection regimes RF were recorded with high tree density followed by PA. Low tree density was observed in SG and PV (Figure 4). The basal area was also found to be significantly different between protection regimes. The high basal area was recorded in SG followed by PA and RF with lowest in PV.

**Figure 4.**
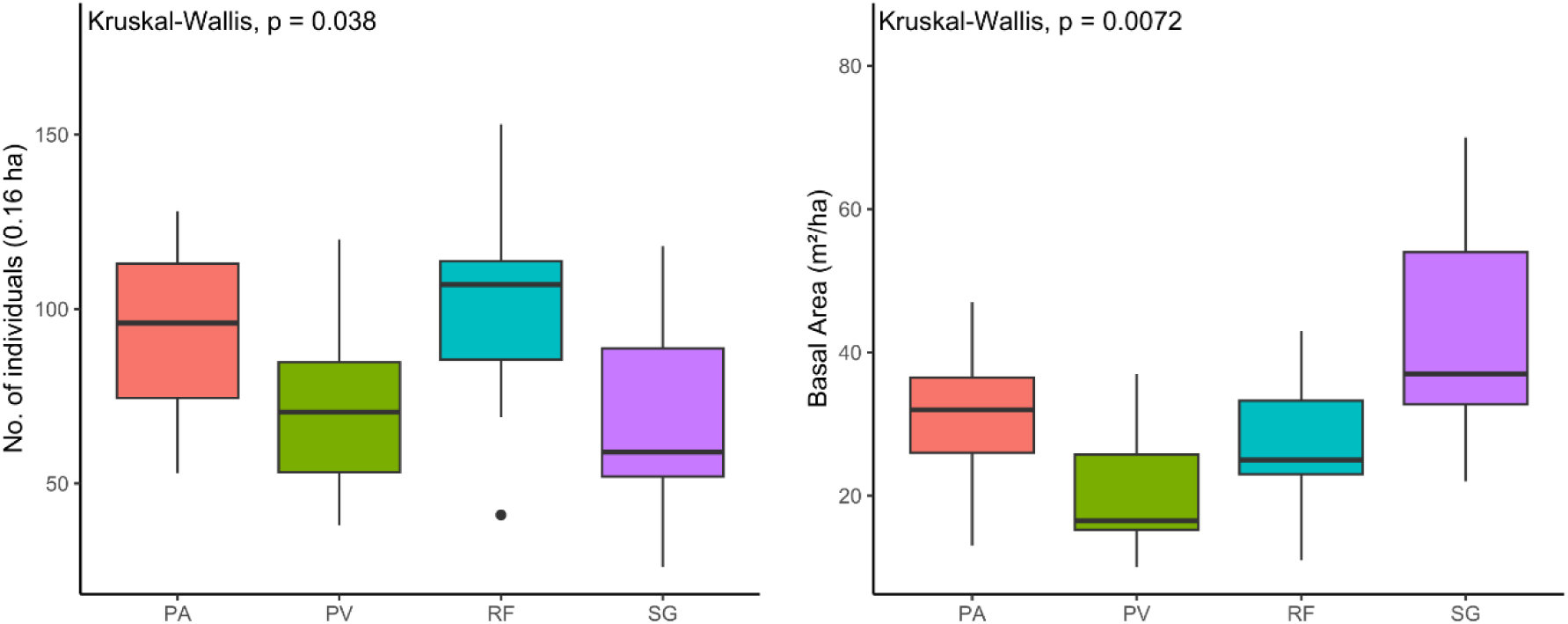
Comparison of number of individuals (0.16 ha^-1^) and basal area (m^2^ ha^-1^) across protection regimes

The analysis of the distribution of number of individuals per girth classes showed that girth classes 30 – 60, 121 – 150 and 181 – 500 were found to be significantly different (Figure 5, Table 2). The girth class 30 – 60 from RF was recorded with higher tree density than other regimes whereas among higher girth classes 121 – 150 and 181 – 500, higher tree density was recorded from SG than other regimes.

**Figure 5.**
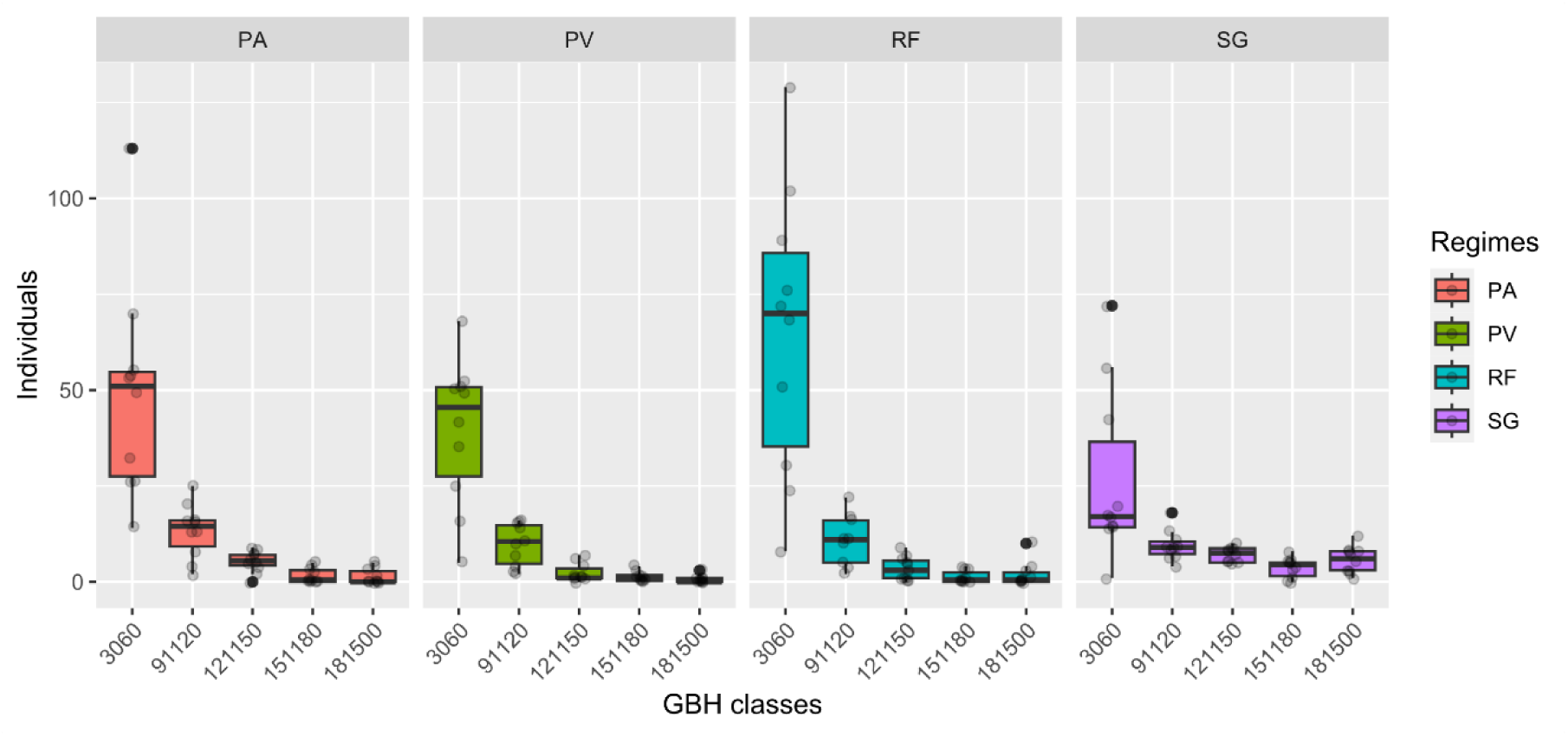
Distribution of number of individuals per girth classes across protection regimes

**Table 2.**
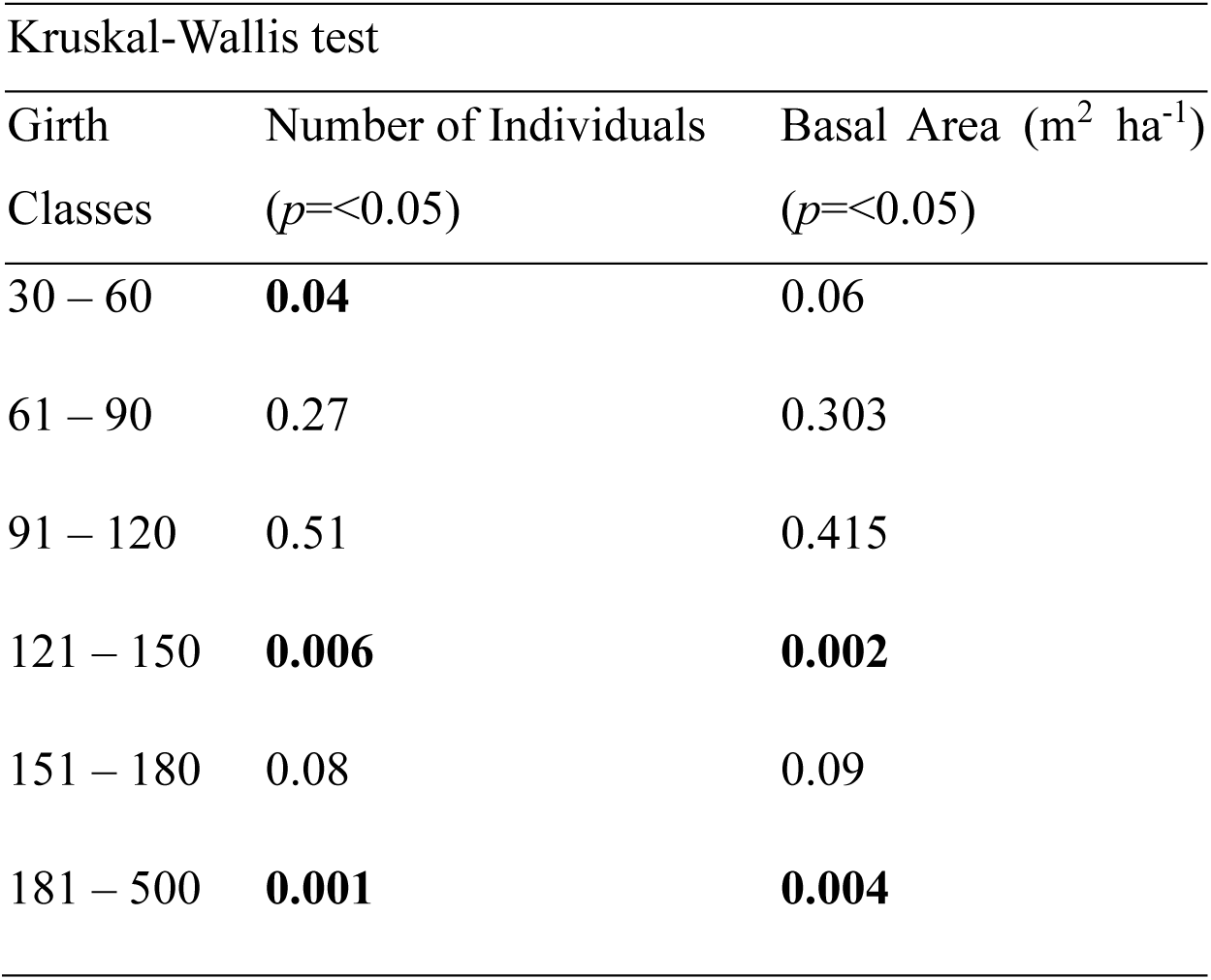
Kruskal-Wallis test score for tree density and basal area in each girth classes.

| Kruskal-Wallis test |  |  |
| --- | --- | --- |
| Girth<br>Classes | Number of Individuals<br>($p < 0.05$) | Basal Area ($\text{m}^2 \text{ ha}^{-1}$)<br>($p < 0.05$) |
| 30 – 60 | **0.04** | 0.06 |
| 61 – 90 | 0.27 | 0.303 |
| 91 – 120 | 0.51 | 0.415 |
| 121 – 150 | **0.006** | **0.002** |
| 151 – 180 | 0.08 | 0.09 |
| 181 – 500 | **0.001** | **0.004** |

The distribution of basal area per girth classes showed significant differences between the classes 121 – 150 and 181 – 500 only (Table 2). It was found that higher basal areas from the same girth classes were recorded from SG than the other regimes. The trees from the 181 – 500 class of PA were also recorded with high basal area followed by SG (Figure 6).

**Figure 6.**
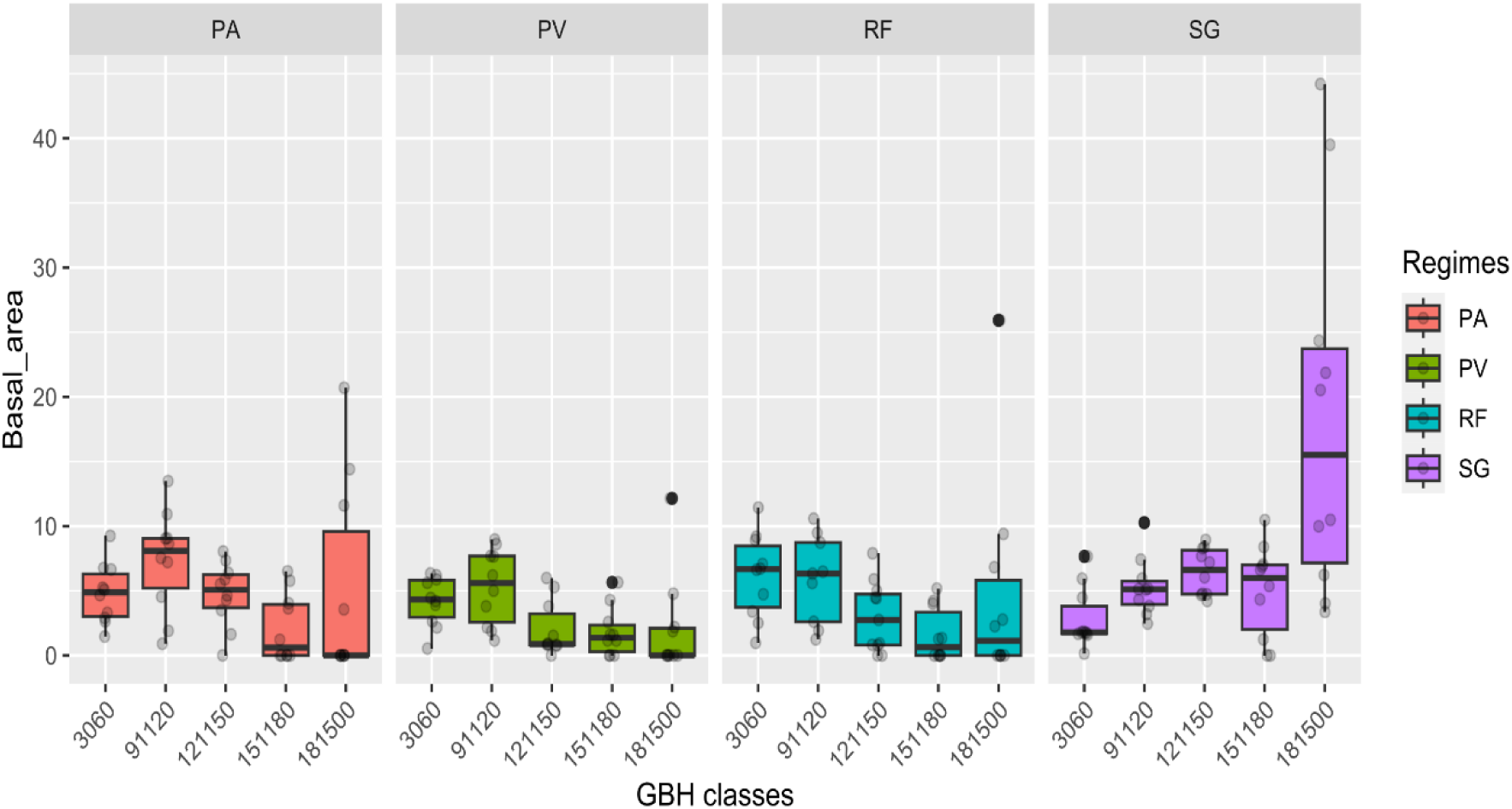
Distribution of basal area per girth classes across protection regimes

## 1.4 Discussion

The forests of the Northern Western Ghats (NWG) have a history of fragmentation and disturbance, currently protected under various regimes. This study evaluates vegetation parameters across four protection regimes, analyzing results in relation to previous research.

### 1.4.1 Species diversity across regimes

The pooled data indicate that species richness and the abundance of evergreen species are higher in RF compared to other protection regimes. However, the average species richness and Shannon index suggest comparable species diversity across the four protection regimes in the Northern Western Ghats (NWG). These findings align with Gardner et al. (2007), who reported minimal variation in species richness among different land use types in Africa, indicating that land use changes may not significantly affect overall species numbers. Similarly, Undaharta et al. (2023) found that both PA and SG in Indonesia exhibited similar levels of species richness and diversity.

In contrast, several studies in India have reported variations in species diversity across protection regimes. For instance, sacred groves in the tropical and subtropical regions of eastern and northeastern India demonstrated higher species richness than protected areas, reserve forests, and non-protected forests (Bdoor, 2016; Pradhan et al., 2019; Rath et al., 2020). A comparative study in the central Western Ghats also found greater species richness in sacred groves than in reserve forests and coffee plantations (Bhagwat et al., 2005b; Ambinakudige & Sathish, 2009). Conversely, community-protected sacred forests in Ghana exhibited lower species richness than protected areas (Opuni-Frimpong et al., 2021). In the current study, species diversity in SGs was comparable to that in PA and RF, consistent with Biswas et al. (2024), who regarded PA and RF as historically disturbed forests and SGs as less disturbed. However, our research found that SGs experienced more disturbance than PA and RF.

### 1.4.2 Community composition across protection regimes

Previous studies have highlighted significant compositional differences across various land use and protection regimes (Ambinakudige & Sathish, 2009; Osuri et al., 2014; Undaharta et al., 2023; Biswas et al., 2024). In contrast, our NMDS analysis indicates a homogeneity in species composition, with approximately 60% dissimilarity reflecting moderate differences among the four regimes. Despite over 50% similarity between SG-PA as well as between PV-RF, the 50% dissimilarity suggests the presence of unique species in each regime. Notably, RF sites like KTRJ and MANO host different species due to their distinct seasonal gradients, with MANO on the crest and KTRJ on the eastern slopes of the NWG. Thus, factors beyond protection level influence species composition. The 50-60% similarity may stem from the underrepresentation of dry deciduous forests in PA and SG, and moderate representation of semi-evergreen to evergreen forests in RF, as most SGs and PAs are located on higher elevations with few deciduous forest representations. Adaptability of forest species and seed dispersal from nearby fragments may also explain species similarities (Bhagwat et al., 2005b; Page et al., 2010).

Among tree species, *Memecylon umbellatum* was the most abundant in PA (26%), RF (10.6%), and SG (19%), while *Garcinia talbottii*, an endemic species, was found primarily in SG (8.9%) and rarely in PA (0.7%) and RF (0.09%). *Terminalia elliptica* was most abundant in PV (16.8%) and second most in RF (9.6%). Unique species recorded only in PA included *Grewia asiatica*, *Streblus aspe*r, and others. PV had 15 unique species, while RF contained 17 unique species, such as *Cochlospermum religiosum* and *Anogeissus latifolia*. SGs also had unique species like *Sageraea laurifolia* and *Artocarpus heterophyllus*.

The rank abundance curve indicated a distribution with many rare species and a few highly abundant ones, reflecting a stable ecosystem with balanced species abundances (Maguran, 2004; Suchiang et al., 2020). The Importance Value Index (IVI) showed moderate changes in dominant species across regimes, suggesting that tree communities are fragments of larger forest areas influenced by varying microhabitats (Hubbell & Foster, 1986; 1992). The dominance of *Memecylon umbellatum* may be due to its status as a shade-intolerant secondary species and early colonizer in degraded areas (Gadgil & Vartak, 1977), thriving in higher elevations during peak shifting cultivation (Ghate et al., 1998). Its wide distribution is aided by seed dispersal by birds and animals (Watve et al., 2003a), though its spread is limited at lower elevations.

*Ficus nervosa*, significant in RF, showed lower abundance despite its contribution to basal area. *Terminalia elliptica* and *T. paniculata* were ecologically significant in PV, likely due to the prevalence of moist to dry deciduous forests at lower elevations. The proportions of evergreen and deciduous species were comparable across regimes, except in PA and PV, where PA sites, located at higher elevations, supported more evergreen trees due to less disturbance (Chandran, 1997; Biswas et al., 2024). PV sites, being highly fragmented and disturbed, were dominated by deciduous tree vegetation.

Seed dispersal plays a crucial role in determining species composition in forest fragments (Butaye et al., 2002). Our study found that zoochory and anemochory methods were comparable across all protection regimes, while autochory was more prevalent in PV and RF, likely due to frequent disturbances creating open spaces (Watve et al., 2003a). Evergreen species predominantly relied on zoochory for dispersal.

### 1.4.3 Comparison of structural aspects

Previous studies have shown that protection regimes and land use types exhibit structural differences in tree density ((i.e. number of individuals ha-1)) and basal area (Gardner et al., 2007; Ambinakudige & Sathish, 2009; Bdoor, 2016; Pradhan et al., 2019; Rath et al., 2019; Biswas et al., 2024). Our analysis similarly revealed variations in tree density and basal area across protection regimes. Specifically, RF and PA had higher tree densities than SG and PV. These findings align with those of Biswas et al. (2024), although SGs in their study had higher densities due to being less disturbed. In contrast, SGs in other regions of India have shown greater tree densities compared to other land uses (Ambinakudige & Sathish, 2009; Bdoor, 2016; Pradhan et al., 2019). Our study, however, recorded lower tree densities in SGs than in other protection regimes, consistent with findings from Opuni-Frimpong et al. (2021) in Ghana. High disturbance levels (CDI) in SG and PV indicate that repeated disturbances hinder tree density, supported by the lower proportion of younger trees (30-60 cm girth class) in these areas. Biswas et al. (2024) also noted reduced tree density in private forests with many cut stems, suggesting that high cutting and logging rates negatively impact young tree populations in SGs and PVs, where average CDIs were 47.75 and 34.5, respectively.

Few studies provide a basis for comparing structural tree vegetation, including those by Kanade et al. (2008) and Joglekar et al. (2015), which reported basal areas in protected forest regions of the NWG ranging from 6.76 to 58.23 m² ha⁻¹. Our study found a basal area of 12.65 to 46.58 m² ha⁻¹ in PAs and 10.54 to 43.18 m² ha⁻¹ in RFs, comparable to Tadwalkar et al. (2020), who reported 27 m² ha⁻¹ in Amboli’s reserve forest. While tree density for younger populations is higher in PAs and RFs, older trees in SGs contribute significantly to basal area. Despite high disturbance and lower abundance in NWG SG, our study recorded a mean basal area of 42.34 m² ha⁻¹, similar to the 36 to 46 m² ha⁻¹ reported by Biswas et al. (2024) in less disturbed SGs. Comparatively, SGs in Meghalaya had a mean basal area of 42.8 m² ha⁻¹ (Mishra et al., 2005), while Tripathi (2003) reported a lower mean of 25.1 m² ha⁻¹. The mean basal area from PVs (20.40 m² ha⁻¹) closely aligns with private forests at lower (24.3 m² ha⁻¹) and higher elevations (11.2 m² ha⁻¹) studied by Biswas et al. (2024). Additionally, a study in Bali, Indonesia, demonstrated a higher basal area in SGs than in PAs (Undaharta et al., 2023).

### 1.4.4 Conservation relevance of protection regimes

This investigation revealed that species richness and composition across protection regimes are not significantly different, indicating that Protected Areas (PA) and Sacred Groves (SG) equally contribute to conservation. Both PA and Reserved Forests (RF) in the Northern Western Ghats (NWG) have a history of disturbances, such as shifting cultivation and forest clearing for settlements (Chandran, 1997; Biswas et al., 2024), and have been under legal protection for 50-100 years. RFs generally support more stable populations than other regimes, contributing to conservation efforts. However, some RF sites near human settlements experience significant interference, including tree cutting, firewood collection, and livestock presence, which may impact the young tree population critical for future carbon mitigation.

Private forests in the region are highly disturbed, showing low numbers of evergreen and endemic trees at lower elevations, highlighting the need for targeted conservation policies (Biswas et al., 2024). The study recommends reintroducing evergreen trees to these sites, as a higher presence of evergreen species correlates positively with basal area, a proxy for biomass and carbon content. Thus, increasing evergreen trees could enhance carbon sequestration, underscoring their importance.

Sacred groves have been protected longer than other regimes (Gadgil & Vartak, 1976) due to cultural beliefs that promote their conservation for ecological services (Ghate et al., 2004). However, urbanization and diminishing religious beliefs have led to increased human interference, resulting in significant loss of sacred groves in the NWG over the past 50 years (Gadgil & Vartak, 1976). Studies indicate substantial losses of sacred groves and above-ground biomass in the central Western Ghats (Bhagwat et al., 2005b; Osuri et al., 2014). Despite these pressures, SGs maintain species diversity comparable to PAs and harbor evergreen, endemic, and unique species, emphasizing the need for conservation efforts focused on these threatened areas. Biswas et al. (2024) highlighted the importance of evergreen species, sacred groves, and reserve forests in lower elevations of NWG, particularly in Sindhudurg. The current study found that many evergreen species are restricted to SGs, and the presence of older trees contributes significantly to basal area, reinforcing the importance of SGs in NWG. The low presence of younger trees warrants further investigation to understand the impact of disturbances on regeneration in SGs and PVs.

## 1.5 Conclusion

In conclusion, while the forests across the four protection regimes—Protected Areas, Community Conserved Forests (Sacred Groves), Reserved Forests, and Privately Owned Forests—exhibit similar species compositions, they significantly differ in structural components such as tree density and basal area. Sacred Groves are characterized by a higher prevalence of older and evergreen trees, whereas Protected Areas and Reserved Forests show a greater representation of younger trees. This highlights the critical need for effective conservation measures in the Northern Western Ghats (NWG).

Without these measures, community-protected forests and endemic species in the region face substantial threats. Disturbances, including the impacts of climate change and potential deforestation driven by changing land use patterns, pose significant risks to the biodiversity of these forests. The consequences of these threats may include a decline in endemic species and an increase in the prevalence of widespread species, disrupting the delicate balance of the ecosystem.

To mitigate these risks and ensure the preservation of biodiversity, urgent and targeted conservation actions are essential. This includes enhancing the protection of Sacred Groves, promoting sustainable land use practices, and addressing the impacts of climate change to safeguard the unique ecological heritage of the NWG.

## Acknowledgements

All authors are thankful to the Director, Agharkar Research Institute for providing facilities for the work. The funding support by Chhatrapati Shahu Maharaj Research, Training and Human Development Institute (SARTHI), Govt. of Maharashtra to BS is acknowledged. Special thanks to APCCF, Forest Department of Maharashtra (letter no: 3353/18-19) for permission to work in the forest areas of the state. ARI inhouse BD-01 funding support to MND and BS is acknowledged.

## Notes

### Competing Interest Statement

The authors have declared no competing interest.

