## SUPPLEMENTARY MATERIAL for "Effect of differential protection regimes on the diversity and composition of woody plants in the Western Ghats"

**Table S1** Family-wise data across protection regimes. The values indicate species richness and the values in the parenthesis indicates number of genera under each family

| **PA** | | **PV** | | **RF** | | **SG** | |
| --- | --- | --- | --- | --- | --- | --- | --- |
| **Families** | **S (G)** | **Families** | **S (G)** | **Families** | **S (G)** | **Families** | **S (G)** |
| Lauraceae | 8 (7) | Leguminosae | 11 (11) | Lauraceae | 6 (6) | Moraceae | 6 (2) |
| Malvaceae | 5 (3) | Rubiaceae | 8 (8) | Rubiaceae | 6 (6) | Lauraceae | 5 (5) |
| Rubiaceae | 5 (5) | Malvaceae | 7 (5) | Anacardiaceae | 5 (4) | Rubiaceae | 5 (5) |
| Anacardiaceae | 4 (4) | Anacardiaceae | 5 (5) | Combretaceae | 5 (2) | Combretaceae | 4 (1) |
| Combretaceae | 4 (1) | Moraceae | 5 (1) | Ebenaceae | 5 (2) | Euphorbiaceae | 4 (4) |
| Euphorbiaceae | 4 (4) | Combretaceae | 4 (1) | Leguminosae | 5 (4) | Leguminosae | 4 (4) |
| Bignoniaceae | 3 (2) | Oleaceae | 3 (3) | Malvaceae | 5 (3) | Anacardiaceae | 3 (3) |
| Moraceae | 3 (2) | Phyllanthaceae | 3 (3) | Moraceae | 5 (2) | Clausiaceae | 3 (2) |
| Myrtaceae | 3 (1) | Rutaceae | 3 (3) | Euphorbiaceae | 4 (4) | Malvaceae | 3 (3) |
| Phyllanthaceae | 3 (3) | Apocynaceae | 2 (2) | Myrtaceae | 4 (2) | Rutaceae | 3 (3) |
| Sapindaceae | 3 (3) | Euphorbiaceae | 2 (2) | Phyllanthaceae | 4 (4) | Sapindaceae | 3 (3) |
| Apocynaceae | 2 (2) | Lamiaceae | 2 (2) | Sapindaceae | 4 (4) | Meliaceae | 2 (2) |
| Clausiaceae | 2 (2) | Lauraceae | 2 (2) | Bursaraceae | 3 (3) | Phyllanthaceae | 2 (2) |
| Ebenaceae | 2 (1) | Sapindaceae | 2 (2) | Clausiaceae | 3 (2) | Sapotaceae | 2 (2) |
| Lamiaceae | 2 (2) | Sapotaceae | 2 (2) | Apocynaceae | 2 (2) | Achariaceae | 1 (1) |
| Leguminosae | 2 (2) | Aracaceae | 1 (1) | Bignoniaceae | 2 (2) | Annonaceae | 1 (1) |
| Meliaceae | 2 (2) | Bignoniaceae | 1 (1) | Lythraceae | 2 (1) | Apocynaceae | 1 (1) |
| Oleaceae | 2 (2) | Clausiaceae | 1 (1) | Rutaceae | 2 (2) | Aracaceae | 1 (1) |
| Rutaceae | 2 (2) | Dilleniaceae | 1 (1) | Annonaceae | 1 (1) | Bignoniaceae | 1 (1) |
| Salicaceae | 2 (2) | Ebenaceae | 1 (1) | Aracaceae | 1 (1) | Bursaraceae | 1 (1) |
| Sapotaceae | 2 (2) | Lecythidaceae | 1 (1) | Bixaceae | 1 (1) | Cannabaceae | 1 (1) |
| Annonaceae | 1 (1) | Longaniaceae | 1 (1) | Boraginaceae | 1 (1) | Celastraceae | 1 (1) |
| Dilleniaceae | 1 (1) | Lythraceae | 1 (1) | Dilleniaceae | 1 (1) | Dilleniaceae | 1 (1) |
| Lecythidaceae | 1 (1) | Melastomataceae | 1 (1) | Icacinaceae | 1 (1) | Lamiaceae | 1 (1) |
| Lythraceae | 1 (1) | Myrtaceae | 1 (1) | Lamiaceae | 1 (1) | Lecythidaceae | 1 (1) |
| Melastomataceae | 1 (1) | Rhamnaceae | 1 (1) | Lecythidaceae | 1 (1) | Lythraceae | 1 (1) |
| Myristicaceae | 1 (1) | Rhizophoraceae | 1 (1) | Melastomataceae | 1 (1) | Melastomataceae | 1 (1) |
| Putranjivaceae | 1 (1) | Symplocaceae | 1 (1) | Meliaceae | 1 (1) | Myristicaceae | 1 (1) |
| Rhizophoraceae | 1 (1) |  |  | Myristicaceae | 1 (1) | Myrtaceae | 1 (1) |
| Symplocaceae | 1 (1) |  |  | Oleaceae | 1 (1) | Oleaceae | 1 (1) |
| Thymelaeaceae | 1 (1) |  |  | Putranjivaceae | 1 (1) | Rhizophoraceae | 1 (1) |
| Ulmaceae | 1 (1) |  |  | Rhizophoraceae | 1 (1) | Salicaceae | 1 (1) |
| Vitaceae | 1 (1) |  |  | Sapotaceae | 1 (1) | Symplocaceae | 1 (1) |
|  |  |  |  | Symplocaceae | 1 (1) | Vitaceae | 1 (1) |
|  |  |  |  | Thymelaeaceae | 1 (1) |  |  |

**Table S2** Test for goodness of fit of different models for observed rank abundance curves across protection regimes (Values in bold indicate fit model)

| **Regimes** | **Model** | **Chi²** | ***p-value*** |
| --- | --- | --- | --- |
| Protected Area (PA) | Geometric | 1851 | 0 |
|  | Log series | 338.3 | 5.10E-44 |
|  | Broken stick | 951.6 | 1.17E-157 |
|  | Log-normal | 1.43 | **0.83** |
| Private Forest (PV) | Geometric | 801.7 | 3.28E-139 |
|  | Log series | 128.2 | 1.87E-06 |
|  | Broken stick | 432.3 | 8.89E-58 |
|  | Log-normal | 4.902 | **0.29** |
| Reserve Forest (RF) | Geometric | 317.8 | 3.88E-37 |
|  | Log series | 21.86 | 1 |
|  | Broken stick | 277.6 | 2.55E-25 |
|  | Log-normal | 5.45 | **0.36** |
| Sacred Grove (SG) | Geometric | 366.3 | 2.14E-52 |
|  | Log series | 46.31 | 0.37 |
|  | Broken stick | 233.7 | 1.44E-23 |
|  | Log-normal | 2.98 | **0.56** |

**Table S3** Importance value index (IVI) for each species across protection regimes (Code details- **Table S5)**

| **Protected area**  **(PA)** | | **Private forests**  **(PV)** | | **Reserve forests**  **(RF)** | | **Sacred groves**  **(SG)** | |
| --- | --- | --- | --- | --- | --- | --- | --- |
| Species | IVI | Species | IVI | Species | IVI | Species | IVI |
| MEMUMB | 52.49 | TERELL | 37.31 | FICNER | 26.90 | MEMUMB | 32.25 |
| OLEDIO | 35.33 | TERPAN | 31.39 | SYZCUM | 23.07 | MANIND | 30.41 |
| SYZCUM | 14.75 | SYZCUM | 24.39 | MEMUMB | 18.87 | SYZCUM | 15.98 |
| DIMLON | 13.63 | OLEDIO | 18.03 | TERELL | 17.33 | OLEDIO | 15.11 |
| FICNER | 10.04 | MEMUMB | 14.19 | OLEDIO | 14.88 | GARTAL | 14.41 |
| XANTOM | 8.37 | BRIRET | 13.22 | TERPAN | 12.91 | IXOBRA | 12.78 |
| MANIND | 8.26 | TECGRA | 11.52 | DIMLON | 9.35 | DIMLON | 11.64 |
| ACTANG | 6.49 | MANIND | 10.31 | XANTOM | 9.34 | TERBEL | 9.11 |
| CARARB | 6.25 | FICREC | 8.03 | FICREC | 7.87 | HOLGRA | 7.39 |
| TERCHE | 6.22 | CARARB | 7.90 | TECGRA | 7.33 | MIMELE | 7.14 |
| IXOBRA | 5.96 | LAGMIC | 7.63 | XYLXYL | 7.16 | STEGUT | 6.74 |
| TERBEL | 5.65 | TERBEL | 6.28 | ATLRAC | 6.38 | XANTOM | 6.60 |
| SCHOLE | 5.20 | ALBCHI | 5.27 | DIOCAN | 6.10 | CARARB | 6.39 |
| TERPAN | 4.93 | BUTMON | 4.98 | CARARB | 6.07 | LAGMIC | 6.28 |
| MIMELE | 4.41 | FICTSJ | 3.78 | LAGMIC | 5.60 | BEIDAL | 5.45 |
| HOLGRA | 4.29 | XANTOM | 3.63 | BRIRET | 4.95 | ELAPAN | 5.33 |
| MYRDAC | 4.19 | CARURN | 3.42 | GRETIL | 4.38 | TERCHE | 5.00 |
| BEIDAL | 3.81 | BOMCEI | 3.31 | DIOSYL | 3.92 | APOLIN | 4.97 |
| ALSSEM | 3.78 | ANAOCC | 3.30 | SCHOLE | 3.52 | DILPEN | 4.76 |
| CATSPI | 3.77 | BAURAC | 3.30 | CARBRA | 3.15 | MAMSUR | 3.69 |
| MACPEL | 3.72 | GRETIL | 3.12 | CATSPI | 3.14 | CHUTAB | 3.50 |
| LAGMIC | 3.71 | CARBRA | 2.87 | WENTHY | 3.10 | FICNER | 3.45 |
| SYZRUB | 3.62 | LANCOR | 2.83 | IXOBRA | 3.04 | FLAMON | 3.34 |
| WENTHY | 3.52 | SPOPIN | 2.72 | LANCOR | 3.03 | FICREC | 3.30 |
| LITSTO | 3.16 | DALLAN | 2.67 | MALPHI | 2.99 | TERPAN | 3.22 |
| GRENER | 3.06 | STRNUX | 2.31 | GLOELL | 2.88 | LEPTET | 3.07 |
| STRTET | 2.83 | IXOBRA | 2.21 | TERBEL | 2.81 | TERELL | 2.99 |
| CRYWIG | 2.80 | GMEARB | 2.19 | ANOLAT | 2.79 | CARURN | 2.99 |
| BRIRET | 2.57 | PTEMAR | 2.04 | DILPEN | 2.73 | ZANRHE | 2.72 |
| TERELL | 2.35 | MADLON | 2.03 | SYMRAC | 2.58 | FICEXA | 2.66 |
| LEPTET | 2.13 | PONPIN | 2.03 | NOTCAS | 2.33 | KNEATT | 2.62 |
| TABALT | 2.11 | XYLXYL | 2.02 | MACPEL | 2.24 | GARIND | 2.52 |
| PONPIN | 2.03 | MEYLAX | 1.95 | HOLGRA | 2.23 | ARTHET | 2.31 |
| LANCOR | 1.98 | PHYEMB | 1.92 | BEIDAL | 2.17 | GMEARB | 2.21 |
| CARBRA | 1.96 | DIOMEL | 1.92 | COCREL | 2.15 | NEOCAS | 2.17 |
| STRASP | 1.90 | MACPEL | 1.81 | APOLIN | 2.11 | BOMCEI | 2.13 |
| DRYVEN | 1.89 | MALPHI | 1.79 | TABALT | 2.09 | LANCOR | 2.07 |
| HETQUA | 1.87 | CANDIC | 1.77 | ACTANG | 2.08 | BRIRET | 2.02 |
| APOLIN | 1.81 | CRYWIG | 1.72 | DIMLAW | 1.88 | FICAMP | 1.97 |
| DILPEN | 1.76 | ACTANG | 1.71 | BUCLAN | 1.86 | ATLRAC | 1.95 |
| GLOELL | 1.76 | MORPUB | 1.65 | MEYLAX | 1.76 | MALPHI | 1.93 |
| FICREC | 1.74 | SYMRAC | 1.50 | CRYWIG | 1.69 | SYMRAC | 1.92 |
| WRITIN | 1.71 | STEURE | 1.39 | SYZGAR | 1.65 | MARIND | 1.75 |
| GREASI | 1.66 | TERCHE | 1.39 | DYSBIN | 1.65 | MURPAN | 1.74 |
| GARTAL | 1.61 | ERINIM | 1.30 | CINVER | 1.62 | MACPEL | 1.71 |
| CLEJAV | 1.56 | MELLUN | 1.26 | HETQUA | 1.57 | MEYLAX | 1.63 |
| MAMSUR | 1.55 | GLOELL | 1.23 | DRYVEN | 1.45 | FICCAL | 1.62 |
| CHIMAL | 1.39 | WRITIN | 1.17 | STEGUT | 1.44 | MACMAC | 1.59 |
| CALTOM | 1.34 | MURPAN | 1.13 | DIOMEL | 1.41 | AGLLAW | 1.49 |
| MACMAC | 1.31 | HOLGRA | 1.12 | DALSIS | 1.40 | NEOCAD | 1.25 |
| LITOLE | 1.29 | SARASO | 1.05 | BOSSER | 1.39 | HYDPEN | 1.21 |
| MITPAR | 1.17 | DILPEN | 1.03 | CANDIC | 1.39 | CANDIC | 1.19 |
| TOOCIL | 1.17 | FICMIC | 1.02 | STEURE | 1.34 | GRETIL | 1.14 |
| MURPAN | 1.16 | CELTIM | 1.00 | HOLARN | 1.34 | SAGLAU | 1.12 |
| MELLUN | 1.16 | KYDCAL | 1.00 | CLEJAV | 1.33 | TAMIND | 1.12 |
| GRETIL | 1.07 | BOMINS | 0.98 | FICAMP | 1.31 | VITALT | 1.09 |
| FALINS | 1.05 | GARIND | 0.96 | GNIGLA | 1.27 | CELTIM | 1.05 |
| STRSP1 | 1.04 | RADXYL | 0.94 | TERCHE | 1.19 | STRTET | 1.04 |
| SYZGAR | 1.04 | ERYSTR | 0.90 | GARPIN | 1.17 | PTEMAR | 1.01 |
| BOMINS | 1.03 | ZIZXYL | 0.90 | EUCGLO | 1.12 | GARPIN | 0.94 |
| GNIGLA | 1.03 | CASFIS | 0.89 | CARURN | 1.03 | SARASO | 0.93 |
| STEGUT | 1.00 | HALCOR | 0.89 | ALSSCH | 0.98 | ALSSCH | 0.90 |
| VITALT | 1.00 | LEPTET | 0.89 | MAMSUR | 0.94 | ACTANG | 0.88 |
| TECGRA | 0.93 | CHIMAL | 0.89 | GARIND | 0.93 | ALSSEM | 0.88 |
| CASFIS | 0.92 | ZANRHE | 0.88 | LAGPAR | 0.92 | CASFIS | 0.87 |
| FLASP1 | 0.91 | DIMLON | 0.88 | DALLAN | 0.88 | FALINS | 0.86 |
| NOTCAS | 0.90 | FICHIS | 0.88 | ALSSEM | 0.87 | CARBRA | 0.85 |
| CHUTAB | 0.83 | WENTHY | 0.88 | NTHNIM | 0.82 | ALLCOB | 0.84 |
| DIOSYL | 0.82 | MITPAR | 0.87 | MEIPAN | 0.80 | CATSPI | 0.84 |
| HOLINT | 0.81 | STEGUT | 0.87 | BOMCEI | 0.78 |  |  |
| SYMRAC | 0.78 | SENCHU | 0.87 | ARTLAK | 0.76 |  |  |
| CASGRA | 0.78 | CATSPI | 0.87 | CANSTR | 0.76 |  |  |
| CINVER | 0.78 | FICEXA | 0.87 | GRENER | 0.73 |  |  |
| MALPHI | 0.78 | HOLPUB | 0.87 | ALBCHI | 0.72 |  |  |
| DIOCAN | 0.78 |  |  | MELLUN | 0.70 |  |  |
| MEYLAX | 0.78 |  |  | GARTAL | 0.70 |  |  |
| MEIPAN | 0.78 |  |  | RADXYL | 0.70 |  |  |
|  |  |  |  | PHYEMB | 0.70 |  |  |
|  |  |  |  | CORDIC | 0.69 |  |  |
|  |  |  |  | FICVIR | 0.69 |  |  |
|  |  |  |  | SYZCAR | 0.69 |  |  |
|  |  |  |  | SAPLAU | 0.69 |  |  |
|  |  |  |  | KNEATT | 0.68 |  |  |
|  |  |  |  | ALLCOB | 0.68 |  |  |
|  |  |  |  | LITSP1 | 0.68 |  |  |
|  |  |  |  | BAURAC | 0.68 |  |  |
|  |  |  |  | HALCOR | 0.68 |  |  |
|  |  |  |  | DIONIG | 0.68 |  |  |
|  |  |  |  | DIOSP1 | 0.68 |  |  |

**Table S4:** Family Index value (FIV) of families and their species richness (S) across protection regimes

| **PA** | | | **PV** | | | **RF** | | | **SG** | | |
| --- | --- | --- | --- | --- | --- | --- | --- | --- | --- | --- | --- |
| **Families** | **FIV** | **S** | **Families** | **FIV** | **S** | **Families** | **FIV** | **S** | **Families** | **FIV** | **S** |
| Melastomataceae | 49.87 | 1 | Combretaceae | 70.05 | 4 | Moraceae | 38.59 | 5 | Anacardiaceae | 35.37 | 3 |
| Oleaceae | 33.44 | 2 | Leguminosae | 32.40 | 11 | Combretaceae | 35.26 | 5 | Melastomataceae | 29.62 | 1 |
| Lauraceae | 23.99 | 8 | Myrtaceae | 21.58 | 1 | Myrtaceae | 25.35 | 4 | Clausiaceae | 21.52 | 3 |
| Sapindaceae | 20.27 | 3 | Anacardiaceae | 20.42 | 5 | Melastomataceae | 17.15 | 1 | Rubiaceae | 20.18 | 6 |
| Myrtaceae | 16.12 | 3 | Moraceae | 17.08 | 5 | Sapindaceae | 14.76 | 4 | Combretaceae | 18.60 | 4 |
| Rubiaceae | 15.82 | 5 | Phyllanthaceae | 14.67 | 3 | Leguminosae | 13.63 | 5 | Moraceae | 17.11 | 6 |
| Combretaceae | 15.20 | 4 | Rubiaceae | 14.28 | 8 | Ebenaceae | 13.28 | 5 | Sapindaceae | 16.47 | 3 |
| Anacardiaceae | 14.73 | 4 | Malvaceae | 14.28 | 7 | Rubiaceae | 13.03 | 6 | Oleaceae | 13.84 | 1 |
| Moraceae | 14.31 | 3 | Lamiaceae | 13.19 | 2 | Oleaceae | 12.59 | 1 | Lauraceae | 13.39 | 5 |
| Sapotaceae | 10.15 | 2 | Oleaceae | 11.95 | 2 | Lauraceae | 10.73 | 6 | Myrtaceae | 12.68 | 1 |
| Malvaceae | 9.74 | 5 | Melastomataceae | 10.36 | 1 | Anacardiaceae | 10.15 | 5 | Sapotaceae | 9.84 | 2 |
| Euphorbiaceae | 7.72 | 4 | Lythraceae | 6.44 | 1 | Phyllanthaceae | 9.45 | 4 | Malvaceae | 9.57 | 3 |
| Bignoniaceae | 7.02 | 3 | Sapotaceae | 5.41 | 2 | Euphorbiaceae | 8.95 | 4 | Phyllanthaceae | 8.96 | 3 |
| Phyllanthaceae | 6.77 | 3 | Rutaceae | 5.24 | 3 | Malvaceae | 8.60 | 5 | Leguminosae | 6.94 | 4 |
| Lecythidaceae | 4.93 | 1 | Lecythidaceae | 4.94 | 1 | Rutaceae | 7.62 | 2 | Rutaceae | 6.64 | 3 |
| Apocynaceae | 4.46 | 2 | Lauraceae | 4.90 | 2 | Sapotaceae | 7.06 | 1 | Meliaceae | 5.82 | 2 |
| Leguminosae | 4.24 | 2 | Euphorbiaceae | 4.30 | 2 | Lamiaceae | 6.18 | 1 | Lecythidaceae | 5.79 | 1 |
| Myristicaceae | 4.18 | 1 | Aracaceae | 3.82 | 1 | Bursaraceae | 4.98 | 3 | Euphorbiaceae | 5.41 | 3 |
| Clausiaceae | 3.80 | 2 | Apocynaceae | 3.32 | 2 | Lythraceae | 4.80 | 2 | Celastraceae | 5.40 | 1 |
| Rutaceae | 3.61 | 2 | Rhizophoraceae | 3.09 | 1 | Lecythidaceae | 4.36 | 1 | Lythraceae | 5.01 | 1 |
| Lamiaceae | 3.57 | 2 | Sapindaceae | 3.08 | 2 | Clausiaceae | 4.23 | 3 | Dilleniaceae | 4.16 | 1 |
| Meliaceae | 3.29 | 2 | Longaniaceae | 2.65 | 1 | Rhizophoraceae | 3.13 | 1 | Salicaceae | 3.41 | 1 |
| Lythraceae | 3.04 | 1 | Symplocaceae | 2.28 | 1 | Apocynaceae | 3.04 | 2 | Myristicaceae | 3.38 | 1 |
| Salicaceae | 2.98 | 2 | Ebenaceae | 1.88 | 1 | Bignoniaceae | 2.81 | 2 | Aracaceae | 2.39 | 1 |
| Ebenaceae | 2.89 | 2 | Dilleniaceae | 1.79 | 1 | Bixaceae | 2.70 | 1 | Lamiaceae | 2.29 | 1 |
| Putranjivaceae | 2.54 | 1 | Cannabaceae | 1.74 | 1 | Symplocaceae | 2.57 | 1 | Symplocaceae | 1.99 | 1 |
| Rhizophoraceae | 1.95 | 1 | Clausiaceae | 1.66 | 1 | Meliaceae | 2.20 | 1 | Achariaceae | 1.96 | 1 |
| Dilleniaceae | 1.75 | 1 | Bignoniaceae | 1.64 | 1 | Dilleniaceae | 2.15 | 1 | Annonaceae | 1.87 | 1 |
| Thymelaeaceae | 1.67 | 1 | Rhamnaceae | 1.57 | 1 | Putranjivaceae | 2.00 | 1 | Vitaceae | 1.84 | 1 |
| Vitaceae | 1.64 | 1 |  |  |  | Thymelaeaceae | 1.83 | 1 | Cannabaceae | 1.81 | 1 |
| Ulmaceae | 1.45 | 1 |  |  |  | Aracaceae | 1.58 | 1 | Bignoniaceae | 1.79 | 1 |
| Symplocaceae | 1.43 | 1 |  |  |  | Icacinaceae | 1.37 | 1 | Bursaraceae | 1.70 | 1 |
| Annonaceae | 1.42 | 1 |  |  |  | Annonaceae | 1.36 | 1 | Apocynaceae | 1.65 | 1 |
|  |  |  |  |  |  | Boraginaceae | 1.25 | 1 | Rhizophoraceae | 1.60 | 1 |
|  |  |  |  |  |  | Myristicaceae | 1.24 | 1 |  |  |  |

**Table S5** Details of code assigned for each taxon during analysis

| **Sr. No.** | **Taxa name** | **Species code** |
| --- | --- | --- |
| 1 | *Actinodaphne lanceolata* | ACTLAN |
| 2 | *Aglaia lawii* | AGLLAW |
| 3 | *Albizia chinensis* | ALBCHI |
| 4 | *Allophylus cobbe* | ALLCOB |
| 5 | *Alseodaphne semecarpifolia* | ALSSEM |
| 6 | *Alstonia scholaris* | ALSSCH |
| 7 | *Anacardium occidentale* | ANAOCC |
| 8 | *Anogeissus latifolia* | ANOLAT |
| 9 | *Aporosa lindleyana* | APOLIN |
| 10 | *Artocarpus heterophyllus* | ARTHET |
| 11 | *Artocarpus lakoocha* | ARTLAK |
| 12 | *Atlantia racemosa* | ATLRAC |
| 13 | *Bauhinia racemosa* | BAURAC |
| 14 | *Beilschmiedia dalzellii* | BEIDAL |
| 15 | *Bombax insignae* | BOMINS |
| 16 | *Bombax ceiba* | BOMCEI |
| 17 | *Boswellia serrata* | BOSSER |
| 18 | *Bridelia retusa* | BRIRET |
| 19 | *Buchanania lanzan* | BUCLAN |
| 20 | *Butea monosperma* | BUTMON |
| 21 | *Callicarpa tomentosa* | CALTOM |
| 22 | *Canarium strictum* | CANSTR |
| 23 | *Canthium dicoccum* | CANDIC |
| 24 | *Carallia brachiata* | CARBRA |
| 25 | *Careya arborea* | CARARB |
| 26 | *Caryota urens* | CARURN |
| 27 | *Casearia graveolens* | CASGRA |
| 28 | *Cassia fistula* | CASFIS |
| 30 | *Celtis timorensis* | CELTIM |
| 31 | *Chionanthus mala-elengi* | CHIMAL |
| 32 | *Chukrasia tabularis* | CHUTAB |
| 33 | *Cinnamomum verum* | CINVER |
| 34 | *Cleidion javanicum* | CLEJAV |
| 35 | *Cochlospermum religiosum* | COCREL |
| 36 | *Cordia dichotoma* | CORDIC |
| 37 | *Cryptocarya wightiana* | CRYWIG |
| 38 | *Dalbergia sissoo* | DALSIS |
| 39 | *Dalbergia lanceolaria* | DALLAN |
| 40 | *Dillenia pentagyna* | DILPEN |
| 41 | *Dimocarpus longan* | DIMLON |
| 42 | *Dimorphocalyx lawinus* | DIMLAW |
| 43 | *Diospyros melanoxylon* | DIOMEL |
| 44 | *Diospyros sylvatica* | DIOSYL |
| 45 | *Diospyros candolleana* | DIOCAN |
| 46 | *Diospyros sp.* | DIOSP1 |
| 47 | *Diospyros nigrescens* | DIONIG |
| 48 | *Drypetes venusta* | DRYVEN |
| 49 | *Dysoxylum binecteriferum* | DYSBIN |
| 50 | *Elaeodendron paniculatum* | ELAPAN |
| 51 | *Erinocarpus nimmonii* | ERINIM |
| 52 | *Erythrina stricta* | ERYSTR |
| 53 | *Eucalyptus globosus* | EUCGLO |
| 54 | *Falconeria insignis* | FALINS |
| 55 | *Ficus racemosa* | FICREC |
| 56 | *Ficus amplisima* | FICAMP |
| 57 | *Ficus tsjakela* | FICTSJ |
| 58 | *Ficus exasperata* | FICEXA |
| 59 | *Ficus nervosa* | FICNER |
| 60 | *Ficus callosa* | FICCAL |
| 61 | *Ficus hispida* | FICHIS |
| 62 | *Ficus microcarpa* | FICMIC |
| 63 | *Ficus virens* | FICVIR |
| 64 | *Flacourtia montana* | FLAMON |
| 65 | *Flacourtia sp.* | FLASP1 |
| 66 | *Garcinia talbotii* | GARTAL |
| 67 | *Garcinia indica* | GARIND |
| 68 | *Garuga pinnata* | GARPIN |
| 69 | *Glochidion ellipticum* | GLOELL |
| 70 | *Gmelina arborea* | GMEARB |
| 71 | *Gnidia glauca* | GNIGLA |
| 72 | *Grewia nervosa* | GRENER |
| 73 | *Grewia asiatica* | GREASI |
| 74 | *Grewia tiliifolia* | GRETIL |
| 75 | *Haldina cordifolia* | HALCOR |
| 76 | *Heterophragma quadriloculare* | HETQUA |
| 77 | *Holarrhena pubescens* | HOLPUB |
| 78 | *Holigarna grahamii* | HOLGRA |
| 79 | *Holigarna arnotiana* | HOLARN |
| 80 | *Holoptelea integrifolia* | HOLINT |
| 81 | *Hydnocarpus pentandrus* | HYDPEN |
| 82 | *Ixora brachiata* | IXOBRA |
| 83 | *Knema attenuata* | KNEATT |
| 84 | *Kydia calycina* | KYDCAL |
| 85 | *Lagerstroemia microcarpa* | LAGMIC |
| 86 | *Lagerstroemia parviflora* | LAGPAR |
| 87 | *Lannea coromandelica* | LANCOR |
| 88 | *Lepisanthes tetraphylla* | LEPTET |
| 89 | *Litsea stocksii* | LITSTO |
| 90 | *Litsea oleoides* | LITOLE |
| 91 | *Litsea sp.* | LITSP1 |
| 92 | *Macaranga peltata* | MACPEL |
| 93 | *Machilus macranthus* | MACMAC |
| 94 | *Madhuca longifolia var latifolia* | MADLON |
| 95 | *Mallotus philippensis* | MALPHI |
| 96 | *Mammea suriga* | MAMSUR |
| 97 | *Mangifera indica* | MANIND |
| 98 | *Margaritaria indica* | MARIND |
| 99 | *Meiogyne pannosa* | MEIPAN |
| 100 | *Melicope lunu-ankenda* | MELLUN |
| 101 | *Memecylon umbellatum* | MEMUMB |
| 102 | *Meyna laxiflora* | MEYLAX |
| 103 | *Mimusops elengi* | MIMELE |
| 104 | *Mitragyna parvifolia* | MITPAR |
| 105 | *Morinda pubescens* | MORPUB |
| 106 | *Murraya paniculata* | MURPAN |
| 107 | *Myristica dactyloides* | MYRDAC |
| 108 | *Neolamarckia cadamba* | NEOCAD |
| 109 | *Neolitsea cassia* | NEOCAS |
| 110 | *Nothopegia castenifolia* | NOTCAS |
| 111 | *Nothopodytes nimmoniana* | NTHNIM |
| 112 | *Olea dioica* | OLEDIO |
| 113 | *Phyllanthus emblica* | PHYEMB |
| 114 | *Pongamia pinnata* | PONPIN |
| 115 | *Pterocarpus marsupium* | PTEMAR |
| 116 | *Radermachera xylocarpa* | RADXYL |
| 117 | *Sageraea laurifolia* | SAGLAU |
| 118 | *Sapindus laurifolius* | SAPLAU |
| 119 | *Saraca asoca* | SARASO |
| 120 | *Schleichera oleosa* | SCHOLE |
| 121 | *Senegalia chundra* | SENCHU |
| 122 | *Spondias pinnata* | SPOPIN |
| 123 | *Sterculia urens* | STEURE |
| 124 | *Sterculia guttata* | STEGUT |
| 125 | *Stereospermum tetragonum* | STRTET |
| 126 | *Stereospermum* | STRSP1 |
| 127 | *Strychnos nux-vomica* | STRNUX |
| 128 | *Symplocos racemosa* | SYMRAC |
| 129 | *Syzygium cumini* | SYZCUM |
| 130 | *Syzygium rubicundum* | SYZRUB |
| 131 | *Syzygium gardneri* | SYZGAR |
| 132 | *Syzygium caryophyllatum* | SYZCAR |
| 133 | *Tabernaemontana alternifolia* | TABALT |
| 134 | *Tamaridus indicus* | TAMIND |
| 135 | *Tectona grandis* | TECGRA |
| 136 | *Terminalia elliptica* | TERELL |
| 137 | *Terminalia chebula* | TERCHE |
| 138 | *Terminalia bellirica* | TERBEL |
| 139 | *Terminalia paniculata* | TERPAN |
| 140 | *Toona cilliata* | TOOCIL |
| 141 | *Streblus asper* | STRASP |
| 142 | *Vitex altissima* | VITALT |
| 143 | *Wendlandia thyrsoidea* | WENTHY |
| 144 | *Wrightia tinctoria* | WRITIN |
| 145 | *Xantolis tomentosa* | XANTOM |
| 146 | *Xylia xylocarpa* | XYLXYL |
| 147 | *Zanthoxylum rhetsa* | ZANRHE |
| 148 | *Ziziphus xylopyrus* | ZIZXYL |

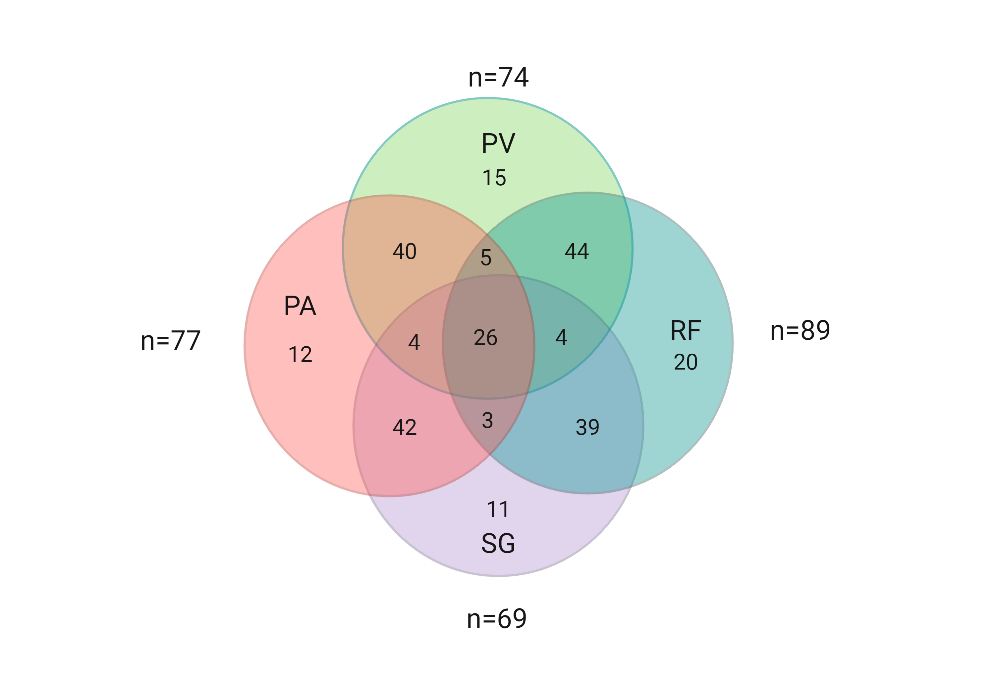

**Figure S1** Comparison of species richness across protection regimes

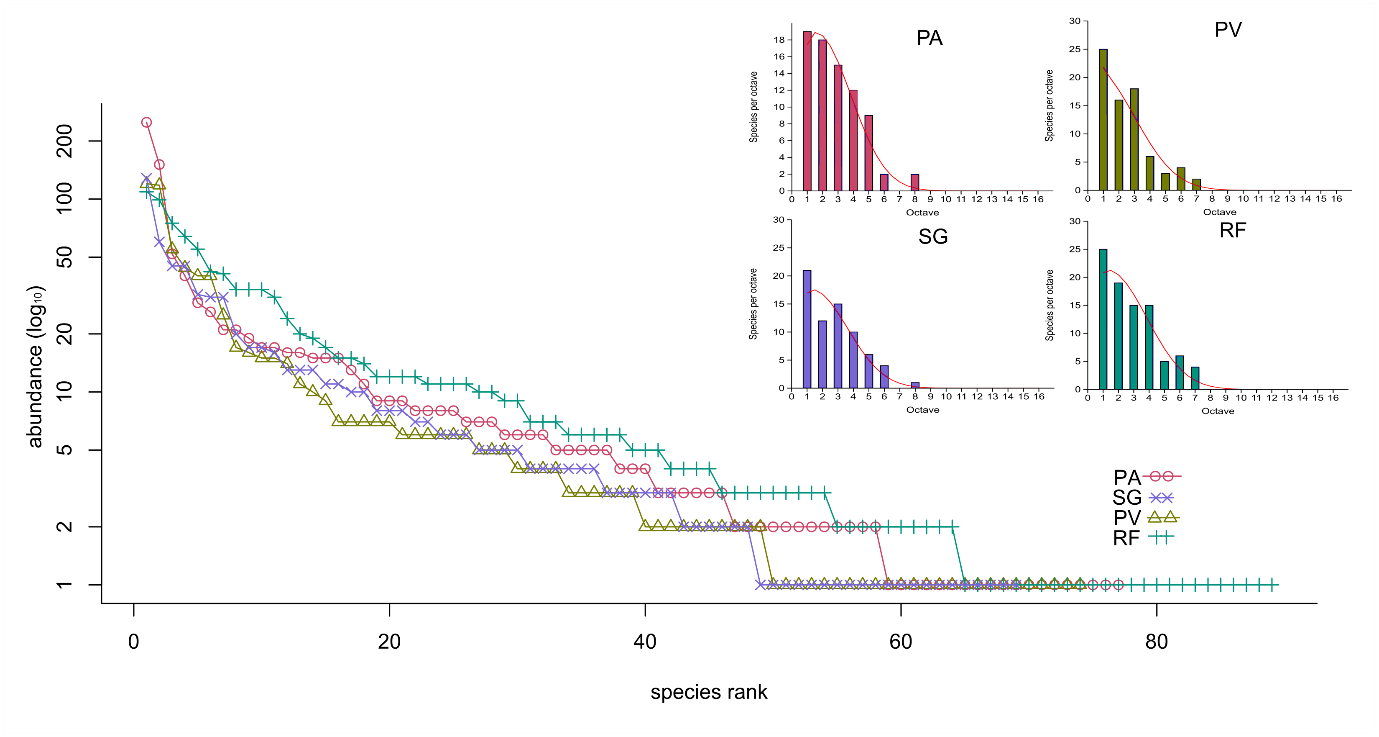

**Figure S2** The log abundance across species rank and log-normal curves for each regime (top right corner)

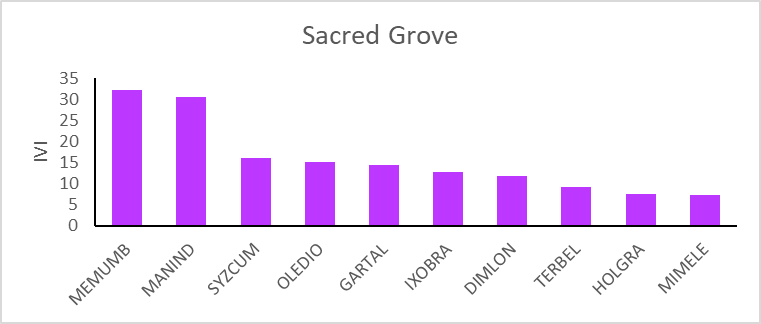

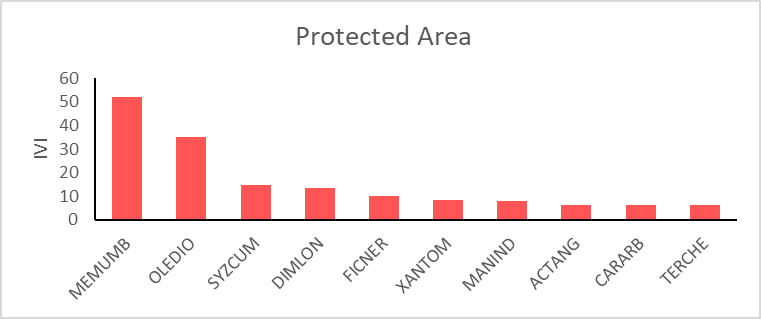

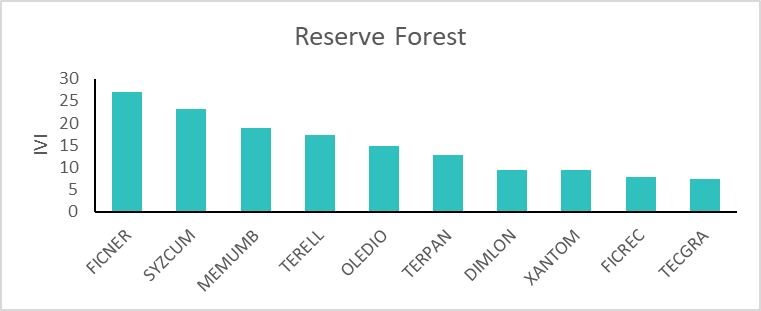

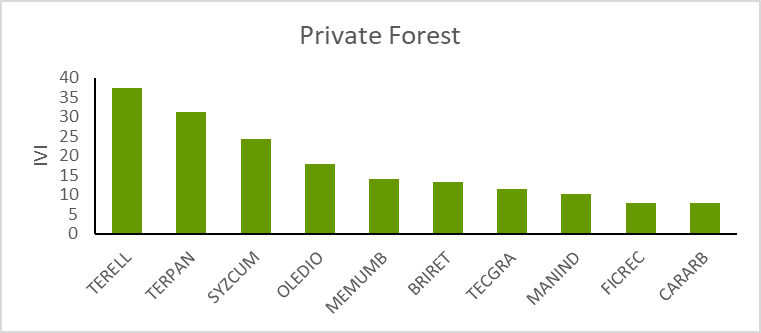

Sacred Grove

IVI

|  | IVI |
| --- | --- |
| MEMUMB | 32.2467 |
| MANIND | 30.4083 |
| SYZCUM | 15.9827 |
| OLEDIO | 15.1127 |
| GARTAL | 14.4072 |
| IXOBRA | 12.7786 |
| DIMLON | 11.6449 |
| TERBEL | 9.10633 |
| HOLGRA | 7.39346 |
| MIMELE | 7.14199 |

Protected Area

IVI

|  | IVI |
| --- | --- |
| MEMUMB | 52.1625 |
| OLEDIO | 35.0723 |
| SYZCUM | 14.6019 |
| DIMLON | 13.5331 |
| FICNER | 9.95962 |
| XANTOM | 8.28438 |
| MANIND | 8.1745 |
| ACTANG | 6.40895 |
| CARARB | 6.17937 |
| TERCHE | 6.14458 |

Reserve Forest

IVI

|  | IVI |
| --- | --- |
| FICNER | 26.8958 |
| SYZCUM | 23.0709 |
| MEMUMB | 18.868 |
| TERELL | 17.3272 |
| OLEDIO | 14.8802 |
| TERPAN | 12.9064 |
| DIMLON | 9.35057 |
| XANTOM | 9.34167 |
| FICREC | 7.86955 |
| TECGRA | 7.33038 |

Private Forest

IVI

|  | IVI |
| --- | --- |
| TERELL | 37.3064 |
| TERPAN | 31.3867 |
| SYZCUM | 24.3904 |
| OLEDIO | 18.0254 |
| MEMUMB | 14.1941 |
| BRIRET | 13.2191 |
| TECGRA | 11.5206 |
| MANIND | 10.3069 |
| FICREC | 8.02719 |
| CARARB | 7.8975 |

**Figure S3** Ten species with the highest Importance Value Index (IVI) per protection regime

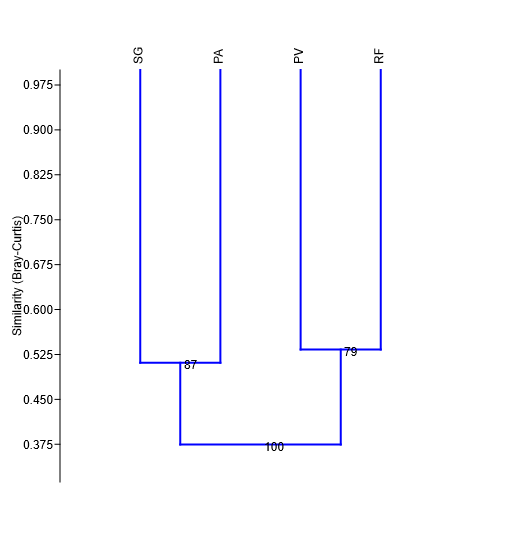

**Figure S4** Cluster analysis showing two distinct clusters based on similarity between protection regimes based on pooled data
